# The sugarcane and sorghum kinomes: insights into evolutionary expansion and diversification

**DOI:** 10.1101/2020.09.15.298612

**Authors:** Alexandre Hild Aono, Ricardo José Gonzaga Pimenta, Ana Letycia Basso Garcia, Fernando Henrique Correr, Guilherme Kenichi Hosaka, Marishani Marin Carrasco, Cláudio Benício Cardoso-Silva, Melina Cristina Mancini, Danilo Augusto Sforça, Lucas Borges dos Santos, James Shiniti Nagai, Luciana Rossini Pinto, Marcos Guimarães de Andrade Landell, Monalisa Sampaio Carneiro, Thiago Willian Balsalobre, Marcos Gonçalves Quiles, Welison Andrade Pereira, Gabriel Rodrigues Alves Margarido, Anete Pereira de Souza

## Abstract

The protein kinase (PK) superfamily is one of the largest superfamilies in plants and is the core regulator of cellular signaling. Even considering this substantial importance, the kinomes of sugarcane and sorghum have not been profiled. Here we identified and profiled the complete kinomes of the polyploid *Saccharum spontaneum* (Ssp) and *Sorghum bicolor* (Sbi), a close diploid relative. The Sbi kinome was composed of 1,210 PKs; for Ssp, we identified 2,919 PKs when disregarding duplications and allelic copies, which were related to 1,345 representative gene models. The Ssp and Sbi PKs were grouped into 20 groups and 120 subfamilies and exhibited high compositional similarities and evolutionary divergences. By utilizing the collinearity between these species, this study offers insights about Sbi and Ssp speciation, PK differentiation and selection. We assessed the PK subfamily expression profiles via RNA-Seq, identifying significant similarities between Sbi and Ssp. Moreover, through coexpression networks, we inferred a core structure of kinase interactions with specific key elements. This study is the first to categorize the allele specificity of a kinome and provides a wide reservoir of molecular and genetic information, enhancing the understanding of the evolutionary history of Sbi and Ssp PKs.

**Highlight:** This study describes the catalog of kinase gene family in *Saccharum spontaneum* and *Sorghum bicolor*, providing a reservoir of molecular features and expression patterns based on RNA-Seq and co-expression networks.

## 1. Introduction

Sugarcane is one of the world’s most important crops, with the highest production quantity and the sixth highest net production value in 2016 (FAO, 2020). For years, this crop has accounted for approximately 80% of the worldwide sugar production (ISO, 2020) and is predicted to account for nearly 40% of the planet’s first-generation biofuel supply in the near future (Lalman *et al*., 2016). However, it is also known for its unprecedented genomic complexity; modern cultivars arose from interspecific crosses between two autopolyploid species, namely, *Saccharum officinarum* (2*n* = 8x = 80, *x* = 10) (D’Hont *et al*., 1998) and *Saccharum spontaneum* (2*n* = 5x =40 to 16x = 128; *x* = 8) (Panje and Babu, 1960). These hybridizations produced large (∼10 Gb) (D’Hont *et al*., 1998), highly polyploid (D’Hont and Glaszmann, 2001) and aneuploid (Sforça *et al*., 2019) genomes. These genomes also contain numerous repetitive elements, mainly retrotransposons, which can account for more than 50% of the total number of sequences (Figueira *et al*., 2012; Kim *et al*., 2013; Mancini *et al*., 2018).

Sugarcane genomic research is severely hampered by this genomic complexity, and for many years depended on resources from a closely related and economically important species: sorghum (*Sorghum bicolor*). *S. bicolor* (Sbi) is a stress-resistant, multifunctional cereal crop that is primarily grown as a staple food in Africa but can also be used for fodder, sugar and biofuel production (Serna-Saldívar *et al*., 2012). *Saccharum* and *Sorghum* belong to the Saccharinae subtribe of the Poaceae family (Clayton, 1987); however, unlike sugarcane, *Sorghum* has not undergone recent polyploidization events (∼96 million years) (Guo *et al*., 2019) and thus has a diploid and much smaller genome that was fully sequenced in 2009 (Paterson *et al*., 2009). Due to both the evolutionary proximity between the two species and the extensive collinearity between their chromosomes, sorghum has historically been considered a diploid model for sugarcane, even before the genome of either species was available (Grivet *et al*., 1996; Grivet and Arruda, 2002).

The superfamily of protein kinases (PKs) comprises the enzymes responsible for catalyzing the reversible phosphorylation of proteins—one of the most widespread posttranslational modifications across all living organisms. PKs act by transferring the terminal phosphate group from adenosine triphosphate (ATP) to the hydroxyl group of a serine, threonine or tyrosine residue in the target protein (Hunter, 1995). These reactions are key events regulating the activity of proteins and protein-protein interactions; therefore, PKs are relevant in many cellular and metabolic processes (Champion *et al*., 2004). In plants, they are involved in the regulation of circadian rhythms and cell cycles, the modulation of various developmental and intracellular processes, and the control of cellular cycles and metabolism (Lehti-Shiu and Shiu, 2012). A recent compilation on stress responses in crops (Hasanuzzaman, 2020) cites numerous reports of the involvement of PKs in plant tolerance to drought, heat and metal toxicity; moreover, many studies have shown that these enzymes play roles in the defense response to herbivores and pathogens (Falco *et al*., 2001; Meng and Zhang, 2013). Several of these responses are predicted to become increasingly relevant in agriculture as a result of climate change; indeed, extreme temperatures and drought are obvious threats from global warming (Dai, 2013; Teixeira *et al*., 2013). Moreover, pest control is also prone to become more challenging with climate instability (Gregory *et al*., 2009). Therefore, the study of molecules and processes associated with both biotic and abiotic stresses is highly relevant to the current setting (Ahuja *et al*., 2010).

Dardick *et al*. (2007) noted that phylogenomic studies are particularly valuable in the analysis of large and conserved gene groups such as PKs because of their ability to form a basis for functional predictions and permit the identification of genes with unique properties, which can in turn allow rational selection of candidates for more detailed studies. The first works on the classification of PKs were based on the conservation and phylogenetic analyses of catalytic domains of eukaryotic proteins (Hanks *et al*., 1988; Hanks and Hunter, 1995). Later studies also considered sequence similarity and domain structure outside the catalytic domains in categorization (Manning *et al*., 2002; Niedner *et al*., 2006). More recently, the availability of low-cost technologies for sequencing whole genomes have allowed the characterization of species’ kinomes, i.e., their entire repertoire of PKs. Compared to the human genome, plant genomes generally contain not only many more PK genes but also atypical kinase families— either exclusive to plant genomes or of prokaryotic origin (Zulawski and Schulze, 2015). This expansion likely resulted from segmental, whole-genome, and tandem duplication events (Hanada *et al*., 2008). *Arabidopsis thaliana* was the first plant to have its kinome compiled (Champion *et al*., 2004), followed by of several other economically important species such as rice, soybean, and grapevine (Dardick *et al*., 2007; Liu *et al*., 2015; Zhu *et al*., 2018b). The kinome of Sbi was compiled shortly after the genome sequencing (Lehti-Shiu and Shiu, 2012).

Several studies have analyzed and characterized kinases in sugarcane. The broadest study is probably the study by Papini-Terzi *et al*. (2005), who identified sequences corresponding to signal transduction components in the sugarcane expressed sequence tag (SUCEST) database (Vettore *et al*., 2003). Although they obtained substantial results considering the limited resources available at the time, these authors reported a relatively low number of PKs (510) in sugarcane. Other studies have indicated that sugarcane PKs are involved in this plant’s development and response to environmental stimuli, such as salt, cold and drought stresses (Carraro *et al*., 2001; Pagariya *et al*., 2012; Li *et al*., 2017). Even more relevant is the compelling evidence that a leucine-rich repeat (LRR) receptor-like kinase is related to sucrose-accumulating sugarcane tissues and genotypes, indicating its involvement in the regulation of sucrose synthesis in mature leaves of this sugar crop (Vicentini *et al*., 2009). However, a more comprehensive identification and characterization of sugarcane PKs has not yet been performed. Recently, a high quality, chromosome-level genome assembly for sugarcane was made available (Zhang *et al*., 2018). The assembly of the genome of the *S. spontaneum* (Ssp) AP85-441 clone (2*n* = 4x = 32) is also allele-defined, i.e., it provides separate sequences of each of the four chromosome copies. The availability of this information-rich reference has since opened a range of possibilities in sugarcane research, such as the detailed characterization of specific groups of genes. Since polyploidy may result in chromosome rearrangements, gene loss and unequal rates of sequence evolution and can favor gene neofunctionalization (Premachandran *et al*., 2011), the sugarcane genome provides fertile ground for related evolutionary and functional studies.

In this context, the main objective of this work was to identify and classify the complete set of PKs present in the Ssp and Sbi genomes. For this purpose, we performed phylogenetic analyses and *in silico* predictions of the properties and subcellular localization of these proteins. Taking advantage of the completeness of the available information, we explored the impact of whole-genome and tandem duplications in the distribution and diversification of the genes encoding PKs in the genomes of these two plants. Finally, we constructed coexpression networks using RNA sequencing (RNA-Seq) to evaluate the expression of PK-encoding genes across different sugarcane and sorghum tissues and genotypes.

## 2. Materials and methods

### 2.1. Kinase identification and domain investigation

All kinase identification and classification procedures were performed for both Sbi and Ssp. The Sbi protein-coding gene sequences and additional files from the Sbi genome (v3.1.1) were obtained from Phytozome v.13 (Goodstein *et al*., 2012). Ssp data were obtained from the AP85-441 genome (Zhang *et al*., 2018) (GenBank accession number: QVOL00000000). The same pipeline was used for both species. All sequences obtained were aligned against the ‘typical’ Pkinase (PF00069) and Pkinase_Tyr (PF07714) families with hidden Markov models (HMMs) retrieved from the Pfam database (El-Gebali *et al*., 2019) using HMMER v.3.3 (Eddy, 1998). An E-value cutoff of 0.1 was used, and we retained only sequences that covered at least 50% of the respective Pkinase domain (Lehti-Shiu and Shiu, 2012). To avoid redundancy, we selected only the longest variant for genes with isoforms. The domain composition of the putative PKs was also investigated via the HMMER web server (Finn *et al*., 2011) and Pfam database. The distribution of PKs across the Sbi and Ssp chromosomes was visualized using MapChart v2.2 software (Voorrips, 2002).

### 2.2. Subfamily classification and phylogenetic analyses

All PKs identified were classified into subfamilies according to HMMs built based on a previous classification and analyses of kinases of 25 plant species (Lehti-Shiu & Shiu, 2012): *Aquilegia coerulea* (Aco), *Arabidopsis lyrata* (Aly), *Arabidopsis thaliana* (Ath), *Brachypodium distachyon* (Bdi), *Carica papaya* (Cpa), *Citrus clementina* (Ccl), *Citrus sinensis* (Csi), *Chlamydomonas reinhardtii* (Cre), *Cucumis sativus* (Csa), *Eucalyptus grandis* (Egr), *Glycine max* (Gma), *Manihot esculenta* (Mes), *Medicago truncatula* (Mtr), *Mimulus guttatus* (Mgu), *Oryza sativa* (Osa), *Populus trichocarpa* (Ptr), *Prunus persica* (Ppe), *Physcomitrella patens* (Ppa), *Ricinus communis* (Rco), *Selaginella moellendorffii* (Smo), *Setaria italica* (Sit), *Vitis vinifera* (Vvi), *Volvox carteri* (Vca), *Zea mays* (Zma), and an earlier version of the Sbi genome, which we called v.1. This classification was confirmed through phylogenetic analyses. The Pkinase domains of the putative PKs were aligned using Muscle v.3.8.31 (Edgar, 2004), and a phylogenetic tree was estimated using a maximum likelihood approach implemented in FastTreeMP v2.1.10 (Price *et al*., 2010) with 1,000 bootstrap replicates using the CIPRES gateway (Miller *et al*., 2010). Different trees were constructed for (I) PKs from Sbi; (II) PKs from Ssp; and (III) PKs from both Sbi and Ssp. The dendrogram visualization and plotting were generated using R statistical software (R Core Team, 2013) with the ggtree (Yu *et al*., 2017) and ggplot2 (Villanueva and Chen, 2019) packages.

### 2.3. Kinase characterization

For each PK identified, the following characteristics were determined: (I) gene chromosomal location and intron number, using GFF files; (II) predicted subcellular localization, with WoLF PSORT (Horton *et al*., 2007), CELLO v.2.5 (Yu *et al*., 2006) and LOCALIZER v.1.0.4 (Sperschneider *et al*., 2017) software; (III) presence of N-terminal signal peptides, using SignalP v.4.1 Server (Petersen *et al*., 2011); (IV) presence of transmembrane domains, using TMHMM v.2.0 Server (Krogh *et al*., 2001); and (V) Gene Ontology (GO) categories (Ashburner *et al*., 2000), using the Blast2GO tool (Conesa *et al*., 2005) with the SWISS-PROT (Bairoch and Apweiler, 2000) and UniProt (UniProt Consortium, 2007) databases. Additionally, for Sbi PKs, alternative splicing (AS) events were investigated using the Plant Alternative Splicing Database (Min, 2013; Min *et al*., 2015). The comparison of these characteristics and calculation of descriptive statistics were performed with R statistical software. Analysis and visualization of GO categories were performed using the REViGO tool (Supek *et al*., 2011) and R (R Core Team, 2013).

### 2.4. Duplication events

To investigate PK duplication events, we used the Multiple Collinearity Scan (MCScanX) toolkit (Wang *et al*., 2012). Tandem duplications were visualized with MapChart v2.2 software (Voorrips, 2002) and segmental events were visualized in circos plots with Circos software (Krzywinski *et al*., 2009). Synonymous substitution (Ks) and nonsynonymous substitution (Ka) rates were also estimated for segmental duplications using MCScanX (Wang *et al*., 2012), and Ks values were used to estimate the date of duplication events: T = Ks/2*λ*, where *λ* is the mean value of the clock-like rates of synonymous substitutions (6.5 ×10−9) (Gaut *et al*., 1996).

### 2.5. RNA-Seq experiments

Data from several RNA-Seq experiments were used to estimate kinase expression. Sbi datasets were retrieved from NCBI’s Sequence Read Archive (SRA) (Leinonen *et al*., 2010) and are described in Supplementary Table S1. Samples from different tissues (pollen, shoots, leaves, microspores, seeds, epidermal tissue, spikelets, roots and internodes) and cultivars (BTX623, BTX642, RTX430, and R07020) were used (Dugas *et al*., 2011; Freeling *et al*., 2015; Makita *et al*., 2015; Kebrom *et al*., 2017; Varoquaux *et al*., 2019). To analyze sugarcane kinase expression, we used novel RNA-Seq datasets described in the following section.

#### 2.5.1. Sugarcane plant material and RNA-Seq

Sugarcane hybrids and *S. officinarum* and Ssp clones were used for expression analyses in sugarcane. Four independent experiments were performed and are detailed in Supplementary Table S2. Experiment 1 was based on root material from the RB867515, RB92579, RB855113, RB855536, SP79-1011, and SP80-3280 hybrid cultivars. This trial was carried out in a greenhouse and used three replicates per cultivar in a completely randomized design. Plants were grown in 18-L plastic pots with a mixture of 20% commercial planting mix and 80% sand. Ninety-five days after planting, we sampled the root material of each plant, avoiding tiller roots.

Experiments 2 and 3 were performed with leaf and culm (internode 1) samples, respectively, from plants grown in the field in Araras, Brazil (22° 18 41.0 S, 47° 23 05.0 W, at an altitude of 611 m). Leaf samples were collected from portions of the top visible dewlap leaves (+1) of six-month-old sugarcane plants in April 2016. We collected the middle section of each leaf, removing the midrib. For culms, samples from the first internode were collected at four time points in 2016: April (synchronous with leaf sampling), June, August and October.

In Experiment 2, we used samples from the SP80-3280, RB72454 and RB855156 hybrid cultivars; TUC71-7 and US85-1008 hybrids; White Transparent and Criolla Rayada *S. officinarum* genotypes; IN84-58, IN84-88, Krakatau and SES205A Ssp genotypes; and IJ76-318 *Saccharum robustum* genotypes. For six genotypes - SP80-3280, RB72454, US85-1008, White Transparent, IN84-58, SES205A - we collected and sequenced three biological replicates, while the others were represented by one biological replicate. All leaf samples were sequenced in two lanes. In Experiment 3, culm samples were collected from the SP80-3280 and R570 hybrid cultivars, F36-819 hybrid, and IN84-58 *S. spontaneum* genotype. Culm samples were sequenced in six lanes.

Experiment 4 was based on samples from the SP80-3280 and IACSP93-3046 hybrid cultivars, Badila De Java *S. officinarum* genotype, and Krakatau Ssp genotype. RNA samples were extracted in triplicate from the top (internode 3) and bottom (internode 8) culms and collected in the field in Ribeirão Preto, Brazil (21° 12 28.7 S, 47° 52 29.1 W) in June 2016.

After collection, samples were immediately frozen in liquid nitrogen and stored at −80°C until processed. Total RNA was extracted from 200 mg of ground roots and 50 mg of ground leaves or culms using an RNeasy Plant Mini Kit (Qiagen, Valencia, CA, United States). We quantified the RNA and verified its integrity in a 2100 BioAnalyzer using a Eukaryote Total RNA Nano Assay (Agilent Technologies). A total of 300 ng of RNA per sample was used to prepare cDNA libraries with a TruSeq Stranded mRNA LT Kit (Illumina, San Diego, USA). All libraries were sequenced on the HiSeq 2500 platform (Illumina, San Diego, USA).

### 2.6. RNA-Seq data processing and coexpression network construction

The quality of the RNA-Seq data was assessed using FastQC software (Andrews, 2010). For read filtering and adapter removal, we used Trimmomatic v.0.39 (Bolger *et al*., 2014). In the Sbi and sugarcane datasets, bases with Phred scores below 20 were removed, and reads shorter than 30 bp were filtered out. In the sugarcane datasets, we also removed the first 12 bases of each read and increased the filter length to 75 bp. For transcript quantification, we used the DNA coding sequences (CDSs) from each species as reference, with k-mers of lengths 31 and 17 for the Ssp and Sbi genomes, respectively, in Salmon v.1.1.0 software (Patro *et al*., 2015). PK expression quantification was evaluated with transcripts per million (TPM) values. Heatmaps visualizing the expression of kinase subfamilies among tissues and cultivars were generated using the R package pheatmap (Kolde and Kolde, 2015) with average TPM values and a complete-linkage hierarchical clustering approach based on Euclidean distances.

Coexpression networks were estimated for PK subfamilies using a minimum Pearson correlation coefficient of 0.6 between PK quantifications across different subfamilies. Network modeling, analysis and visualization were performed using the R package igraph (Csardi and Nepusz, 2006). To assess the Ssp and Sbi network structures and subfamily characteristics within the networks, hub scores for each subfamily were calculated considering Kleinberg’s hub centrality scores (Kleinberg, 1999), edge betweenness values estimated by the number of geodesics passing through the edge (Brandes, 2001), and communities defined using a propagating label approach (Raghavan *et al*., 2007).

## 3. Results

### 3.1. Genome-wide identification of PKs in sugarcane and sorghum

All Ssp and Sbi protein sequences available were aligned against kinase domains using HMMER, and 3,729 (Ssp) and 1,910 (Sbi) different sequences showed significant correspondence with Pkinase families (minimum E-value of 0.1). To avoid redundancies in this set, we removed Sbi isoforms by using its GFF file, resulting in 1,276 sequences. Additionally, the kinase domain coverage of all Ssp and Sbi alignments were evaluated; 810 (Ssp) and 66 (Sbi) sequences did not have a minimum domain coverage of 50% and were therefore discarded, as they likely represented atypical kinases or pseudogenes (Lehti-Shiu and Shiu, 2012; Liu *et al*., 2015). Ultimately, we identified 2,919 putative Ssp and 1,210 putative Sbi PKs. Supplementary Tables S3 and S4 show the discrimination of the kinase domain correspondence for the selected sequences; the data indicate that some PKs (228 Ssp and 49 Sbi PKs) contained multiple kinase domains.

Genome-wide identification of Ssp PKs was performed without prior knowledge of allelic relationships among genes; however, due to the allele specificity of Ssp PKs, we also evaluated their gene model (GM) organization as defined by Zhang *et al*. (2018). These authors associated different sets of allele copies, paralog and tandem duplications to only one representative GM. The 2,919 Ssp PKs corresponded to 1,345 different GMs, and the number of Ssp PKs was only ∼5% higher than that of Sbi PKs, which did not include allele differences. By analyzing the GM description file (Zhang *et al*., 2018), we identified 3,717 different gene configurations for the 1,345 selected GMs, exceeding the number of detected kinases (2,919). However, these divergences in quantity were related only to tandem and paralogous duplications, and the number of allele copies (2,575) was identical in both analyses.

The Ssp and Sbi PKs were further classified into groups and subfamilies using HMMs built based on the kinase sequences of 25 plant species identified by Lehti-Shiu & Shiu (2012). PKs were classified into the subfamily with the top-scoring HMM correspondence. This process resulted in the identification of 119 kinase subfamilies in Ssp and 120 in Sbi (Supplementary Tables S5 and S6), corresponding to 20 different groups. This classification was confirmed by three different phylogenetic trees (Supplementary Figs. S1-S3) estimated based on Sbi PKs, Ssp PKs and all PKs from the two species. In the dendrogram, only 7 sequences in Ssp and 2 in Sbi did not cluster with any other kinase subfamily. These unclassified PKs were included in an “Unknown” category and considered probable novel gene kinase subfamilies. Comparison of the Ssp GM and Sbi PKs revealed that the number and relative proportion of proteins in each group was similar (Supplementary Table S7) with 40% of subfamilies’ quantities having the same values for Ssp and Sbi.

Overall, the number of PKs in each subfamily was low. The mean number of PKs per subfamily was 10 in Sbi (median, 3), and 25 in Ssp (median, 4). The most abundant group in both species was the receptor-like kinase (RLK)-Pelle group, accounting for ∼70% of the PKs, followed by the calcium- and calmodulin-regulated kinase (CAMK); cyclin-dependent kinase, mitogen-activated protein kinase, glycogen synthase kinase and cyclin-dependent kinase-like kinase (CMGC); tyrosine kinase-like kinase (TKL); serine/threonine kinase (STE); and cyclic AMP-dependent protein kinase (cAPK), cGMP-dependent protein kinase, and lipid signaling kinase families (AGC); and casein kinase 1 (CK1) groups. All other groups contained less than 1% of the total number of PKs. The clear separation and high abundance of the RLK group can be seen clearly in Fig. 1. These subfamily abundances were similar for Ssp and Sbi, and only the pancreatic eukaryotic initiation factor-2alpha kinase (PEK_PEK) subfamily was exclusive to Sbi. The absolute counts ranged from 1 to 189 in Ssp GMs and from 1 to 133 in Sbi; the most abundant subfamilies in these species were RLK-Pelle_DLSV (14.04% in Ssp and 10.99% in Sbi), RLK-Pelle_WAK (4.38% in Ssp and 6.12% in Sbi), RLK-Pelle_L-LEC (6.84% in Ssp and 5.7% in Sbi), RLK-Pelle_SD-2b (6.39% in Ssp and 4.96% in Sbi), RLK-Pelle_LRR-XII-1 (2.67% in Ssp and 4.79% in Sbi), RLK-Pelle_LRR-XI-1 (4.68% in Ssp and 4.21% in Sbi) and CAMK_CDPK (3.27% in Ssp and 3.22% in Sbi).

**Fig. 1:**
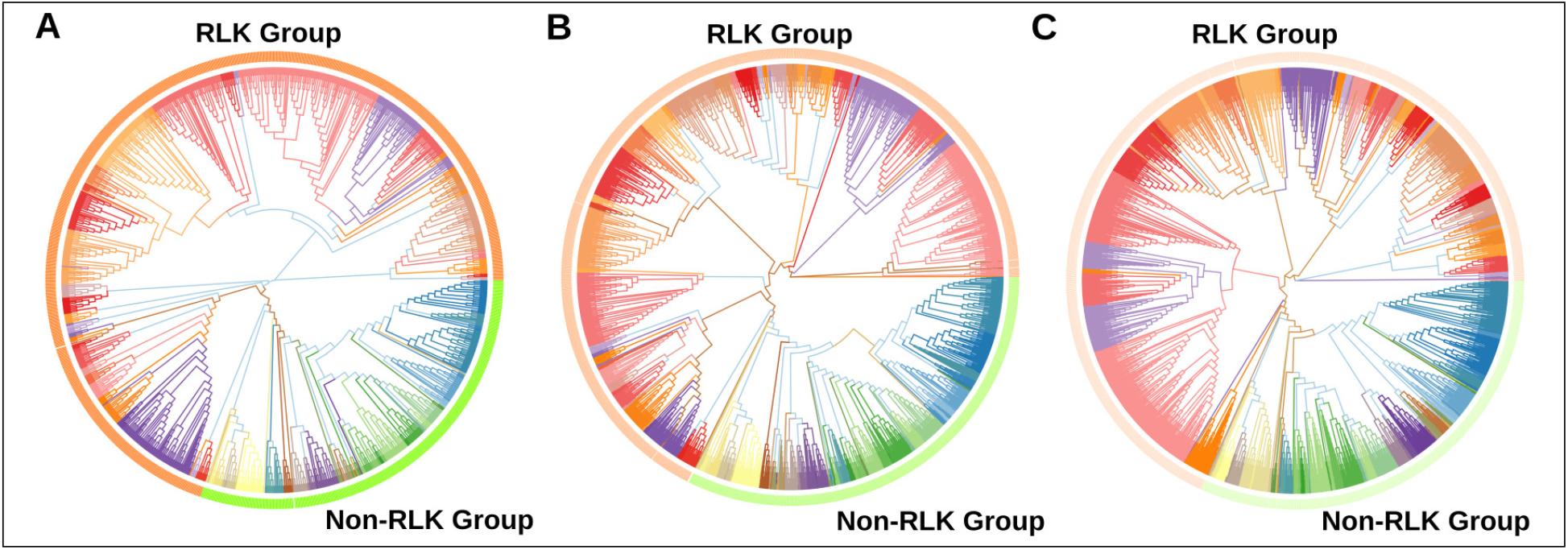
Phylogenetic analyses of putative protein kinases (PKs) identified in the *Saccharum spontaneum* (Ssp) and *Sorghum bicolor* (Sbi) genomes. (A) Phylogenetic tree of the 1,210 Sbi PKs organized in 120 subfamilies represented by different colors. (B) Phylogenetic tree of the 2,919 Ssp PKs organized in 119 subfamilies. (C) Phylogenetic tree of PKs in both Sbi and Ssp.

Additionally, we compared the identified subfamily quantities to 25 other plant species included in the study of Lehti-Shiu & Shiu (2012). The heatmap (Supplementary Fig. S4) visualizing the similarities in the numbers of PKs indicated a closeness between the Ssp and Sbi kinase compositions; however, both exhibited closer relationships with other species than with each other. The dendrogram constructed based on the columns (plant species) enabled the identification of the species most similar to Sbi and Ssp in terms of PK quantities. Sbi was found to belong to a cohesive clade with Zma, Bdi, and Sbi v.1; Ssp belonged to a clade with Sit and Osa. Interestingly, even though these groups were separated by other species, together, these two clusters corresponded to all of the monocotyledon species used in this comparison, and the other clusters corresponded to dicotyledon species, bryophytes and green algae. Considering these two clusters containing Sbi and Ssp, 26 subfamilies were not represented by PKs, corroborating the correspondence of the PK quantities between these species.

### 3.2. Characterization of PKs

Ssp and Sbi PKs were distributed across all Ssp and Sbi chromosomes and alleles (Fig. 2A and B). We found 1,209 PKs among all 10 Sbi chromosomes and 1 PK in a separated scaffold (Supplementary Table S8). The Sbi PK quantities ranged from 67 (5.54%) on chromosome 7 to 184 (15.21%) on chromosome 3. In Ssp (Supplementary Table S9), the PK quantities across allelic configurations were similar (762 in A, 748 in B, 675 in C, and 734 in D), and in all configurations, chromosome 2 had the most and the chromosome 6 had the fewest PKs. The accumulation of PKs was generally consistent with an increase in the chromosomal length.

**Fig. 2:**
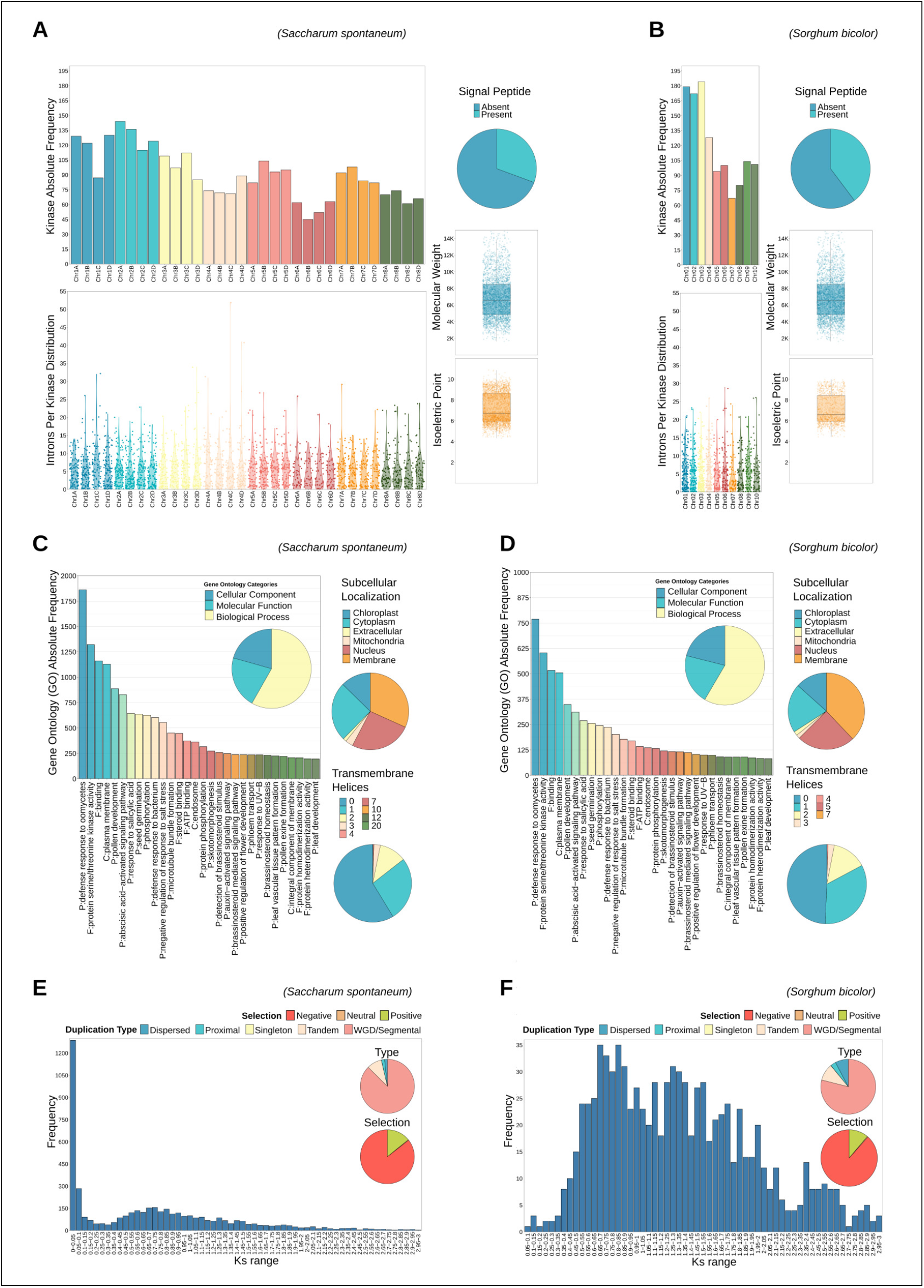
Descriptive analysis of kinase characteristics in *Saccharum spontaneum* (Ssp) (A, C, and E) and *Sorghum bicolor* (Sbi) (B, D, and F): chromosomal distribution, intron length and chromosomal occurrence, presence of signal peptides, molecular weights (MWs), isoelectric points (pIs), Gene Ontology (GO) terms, subcellular localization, presence of transmembrane helices and duplication events.

The intron distribution differed between Ssp and Sbi PKs (Supplementary Tables S10 and S11) and did not exhibit a clear distribution pattern on a specific chromosome (Fig. 2A and B). A large number of PKs were intronless (154 in Sbi and 329 in Ssp). Additionally, all identified PKs were analyzed against the Pfam database to retrieve related nonkinase domains. In Sbi, we identified 70 additional domains (Supplementary Table S12) distributed across 662 PKs (Supplementary Table S13). Interestingly, none of these additional domains were found in the 49 PKs containing multiple kinase domains (Supplementary Table S14). In Ssp, we identified 168 additional domains (Supplementary Table S15) across 1,423 PKs (Supplementary Table S16). The 228 Ssp PKs with multiple kinase domains (Supplementary Table S17) also did not present nonkinase domains. These additional domains were similar in Sbi and Ssp PKs (60 domains in common). The 5 most abundant domains in Sbi were LRRs, with 8 in 193 PKs; leucine-rich repeat N-terminal domains (LRRNTs), with 2 in 192 PKs, D-mannose-binding lectin (B-lectin) domains, in 83 PKs; wall-associated receptor kinase (WAK) galacturonan-binding (GUB) domains, in 72 PKs; and S-locus glycoprotein domain (S-locus-glycop) domains, in 71 PKs. In Ssp, the 5 most abundant domains were LRRNTs, with 2 in 390 PKs; LRRs, with 8 in 351 PKs; B-lectin domains, in 218 PKs; S-locus-glycop domains, in 185 PKs; and PAN-like domains (PAN_2), in 182 PKs. Four of the five domains had the same abundance ranking in both species.

A full GO annotation of Sbi and Ssp PKs was performed with Blast2GO (Supplementary Tables S18 and S19). In Sbi, we found 1,581 different GO terms related to 18,320 correspondences among the PKs. These terms were separated into 3,857 (21.05%) terms related to the cellular component GO category, 3,752 (20.48%) to the molecular function category, and 10,711 (58.47%) to the biological process category. In Ssp, we found more categories (1,875) and more correspondences (44,582) due to the larger size of the Ssp kinome. However, the proportion of GO terms was similar: 9,193 (20.62%) in the cellular component GO category, 9,429 (21.15%) in the molecular function GO category, and 25,960 (58.23%) in the biological process GO category. This clear similarity can be observed in the GO analysis pie charts in Fig. 2C and D. The 30 most abundant GO terms in Sbi and Ssp (with 26 categories in common) are also shown. Due to the clear comprehensiveness of GO categories related to biological processes, an additional analysis was performed using these Sbi and Ssp GO terms. Using REViGO software, treemaps were generated to summarize these categories based on semantic similarities (Supplementary Fig. S5A and B); the most abundant biological processes were related to protein phosphorylation, defense response and cellular development.

For Sbi PKs, we investigated the possible occurrence of alternative splicing using the Plant Alternative Splicing Database. One hundred Sbi kinase genes were found to undergo associated alternative splicing events, and GO analysis of the most frequent biological processes associated with these genes (Supplementary Fig. S5C) showed changes in the most frequent categories considering the entire dataset of PK-related GO terms. The most frequent category organization was defense response, which was the third most frequent in the entire set of Sbi PK GO terms. In addition, programmed cell death was included as a category organization instead of cell growth.

We also explored the presence of signal peptides and transmembrane helices in the PKs and investigated their estimated molecular weights (MWs), theoretical isoelectric points (pIs), and subcellular localization (Supplementary Tables S20 and 21). Among the Sbi PKs, ∼40% were predicted to contain signal peptides (Fig. 2A), in contrast with ∼30% of Ssp PKs (Fig. 2B). The MW and IP distributions of Sbi and Ssp PKs are shown in Fig. 2A and B, respectively. Most Ssp PKs (∼59%) did not contain transmembrane helices, while 50% of Sbi PKs did not. The remaining sequences in both Sbi and Ssp PKs contained many (between 1 and 3) transmembrane helices (Fig. 2C and D). To predict the subcellular localization of PKs, we used three different software packages (WoLF PSORT, CELLO and LOCALIZER). The results indicated high divergence among these methods; thus, we considered only the predictions identified by a consensus of at least two of the three tools used. The localization of 1,425 Ssp and 616 Sbi PKs was predicted. The PKs were classified as localized in the chloroplast, cytoplasmic, extracellular, mitochondrial, nuclear or membrane regions (Fig. 2C and D). The most frequently identified localization was the membrane, as also indicated by the high frequency of the plasma membrane GO term.

The attributes of the PKs are summarized at the kinase subfamily level in Supplementary Table S22 for Sbi and in Supplementary Table S23 for Ssp. To characterize kinase subfamily gene structures, we first calculated the mean quantity of introns per kinase in each subfamily and then determined the standard deviation and the coefficient of variation. As already shown (Supplementary Tables S5 and S6), several subfamilies contained only one representative gene (30 in Sbi and 33 in Ssp). In Ssp, some of these subfamilies with one GM had high intronic divergences in gene allelic copies (with coefficients of variation ranging from 0 to ∼141%). Considering only the subfamilies with more than one member, increased coefficients of variation were observed (ranging from 0 to ∼241%), corresponding to significant discrepancies in gene organization within kinase subfamilies. By filtering the subfamilies with a maximum coefficient of variation of 20% and at least 2 PKs, we identified only 37 Sbi and 12 Ssp subfamilies with a more cohesive structure, but most of these included only a few PKs. The five subfamilies among these structurally organized groups with the highest number of PKs were RLK-Pelle_LRR-I-2, TKL_CTR1-DRK-2, NEK, CK1_CK1-Pl and PEK_GCN2 in Ssp and RLK-Pelle_LRR-II, RLK-Pelle_LRR-I-1, RLK-Pelle_LRR-V, RLK-Pelle_RLCK-VIII, and RLK-Pelle_RLCK-V in Sbi. Interestingly, the highest intron numbers were also observed in members of subfamilies belonging to RLK-Pelle groups, with the exception of PEK_GCN2 in Sbi.

Protein properties across kinase subfamilies were also summarized and did not exhibit considerable differences. Based on a maximum coefficient of variation of 20%, 13 subfamilies in Sbi and 15 in Ssp had considerable variations in the IP. The MW exhibited higher variability in Ssp than in Sbi (66 subfamilies with more diverse values, in contrast to 20 in Sbi). Regarding the presence of signal peptides, all PKs in only 18 Sbi PK subfamilies (6 of which contained only one PK) contained these subsequences; the subfamilies with the most members were RLK-Pelle_LRR-V (12 members) and RLK-Pelle_WAK_LRK10L-1 (7 members). In Ssp, all PKs in only 8 subfamilies contained signal peptides, with the inositol-requiring kinase 1 (IRE1) and RLK-Pelle_RLCK-X subfamilies each containing 5 members. Similarly, these highlighted subfamilies also contained transmembrane helices across their kinomes.

To complement the protein properties observed in kinase subfamilies, the domain composition was described (Supplementary Tables S24 (Sbi) and S25 (Ssp)). Interestingly, the AGC_RSK-2 subfamily had the highest number of PKs with multiple kinase domains in both Sbi (19 PKs) and Ssp (20 PKs). Furthermore, we investigated the percentage of multikinase domain-containing proteins among the PKs in each subfamily (Supplementary Tables S22 and S23). The highest percentage (100%) was observed in the AGC_NDR and CMGC_SRPK subfamilies in Sbi and in the CMGC_SRPK, CMGC_CDK-CCRK subfamilies in Ssp. Even though the AGC_NDR subfamily did not contain all of the proteins with multiple kinase domains in Ssp, 10 of the 15 (∼66%) had this characteristic. In general, the same domains were observed in Sbi and Ssp, as already described. Across subfamilies, the 10 most abundant protein domains were almost the same in Sbi and Ssp and comprised the LRRNT_2, LRR_8, LRR_1, LRR_6, LRR_4, B_lectin, S-locus-glycop, GUB_WAK_bind, salt stress response, and antifungal domains. The 10 subfamilies with more varied domains belonged to the RLK-Pelle group in Sbi. However, in Ssp, the CMGC_CDK group was the most domain-diverse subfamily.

### 3.3. Kinase duplication events in sugarcane and sorghum

Gene duplications in Sbi and Ssp kinases were investigated using MCScanX. We identified numerous kinase genes (1,165 in Sbi and 2,919 in Ssp) with an origin associated with dispersed (7.73% in Sbi and 1.68% in Ssp), proximal (3.18% in Sbi and 1.88% in Ssp), tandem (10.04% in Sbi and 8.94% in Ssp) and segmental duplications (78.97% in Sbi and 87.43% in Ssp). These classifications are described in Supplementary Tables S26 and S27. Ssp PKs with origins related to tandem duplications were differentially distributed across all allele copies on chromosomes (ranging from 2 events on Chr4-B to 16 events on Chr3-A). The mean value per allele was 8.16, with the highest concentration in allele copies on chromosome 3. A visual map of all Ssp PKs organized in tandem was constructed using chromosomal representations according to their physical location retained in the GFF file and were colored according to kinase subfamilies (Fig. 3B). Tandemly organized Sbi PKs were also visualized (Fig. 4B). All Sbi chromosomes contained PKs with origins associated with tandem duplications. Chromosome 4 contained only 1 PK with a tandem duplication-associated origin, and chromosomes 2 and 3 had the most such PKs (26 PKs on both). By analyzing the tandemly duplicated PKs within subfamilies, we found 19 subfamilies containing PKs that originated by tandem duplication. The highest percentages of such Sbi PKs were found in the RLK-Pelle_RLCK-Os (80%), RLK-Pelle_LRR-I-1 (37.5%), CMGC_CDKL-Os (34.78%), RLK-Pelle_LRK10L-2 (34.48%), and CMGC_CK2 33.33%) subfamilies. In Ssp, 64 subfamilies had tandemly duplicated PKs, and the 5 subfamilies with the highest percentages were RLK-Pelle_RKF3 (100%), RLK-Pelle_LRR-VIII-1 (37.5%), CAMK_CAMK1-DCAMKL (33.33%), RLK-Pelle_LRR-XIIIb (30%), and TKL_Gdt (28.57%).

**Fig. 3:**
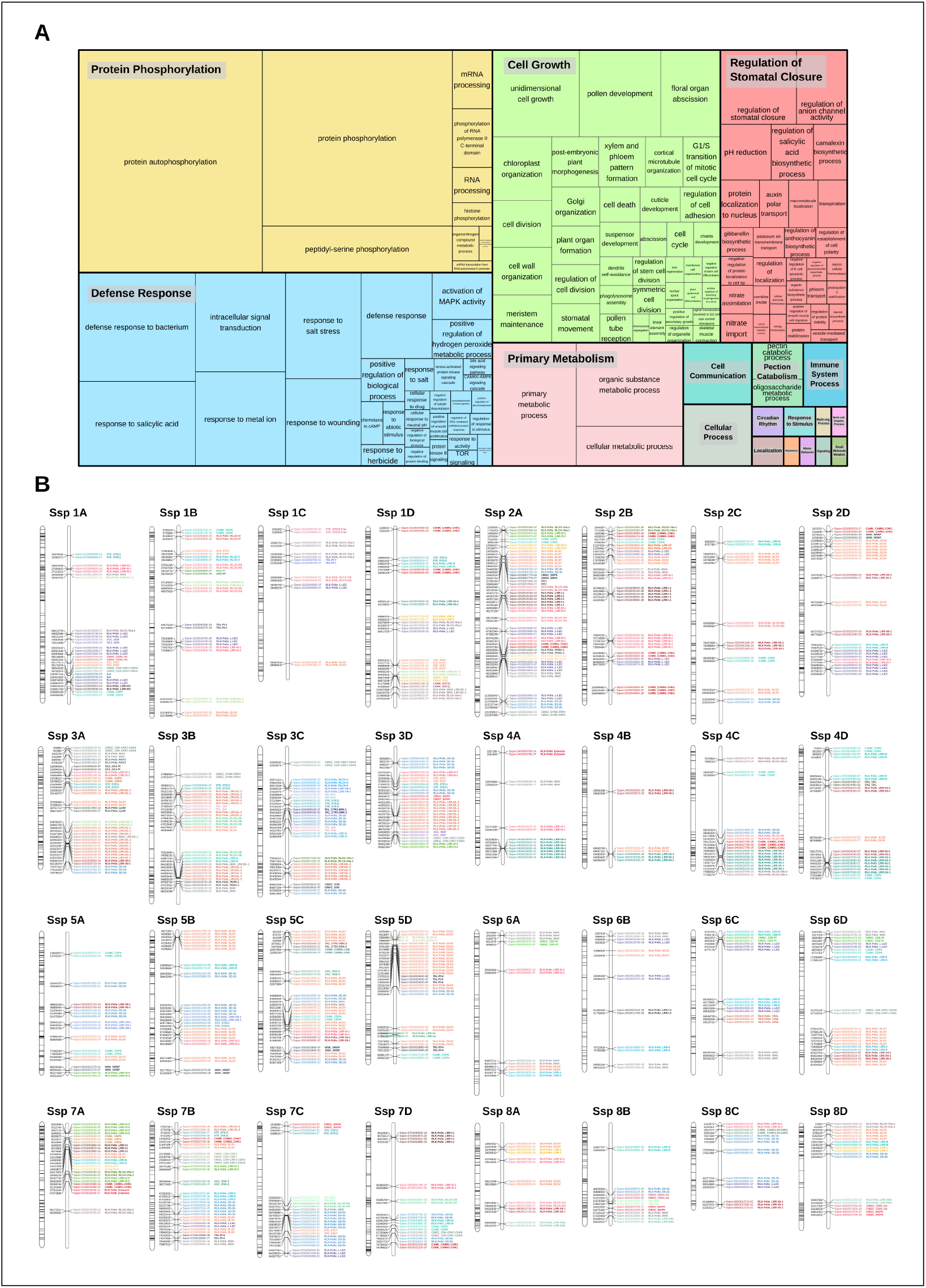
(A) Gene Ontology (GO) categories (biological processes) related to tandemly duplicated kinases in *Saccharum spontaneum* (Ssp). The size of the subdivisions within the blocks represents the abundance of that category in this set of kinases. The colors are related to the similarity to a representative GO annotation for the group. (B) Kinase distribution along Ssp chromosomes. For each chromosome, all genes with kinase domains are indicated on the left, and only the tandemly organized kinases are indicated on the right, colored and labeled according to the subfamily classification.

**Fig. 4:**
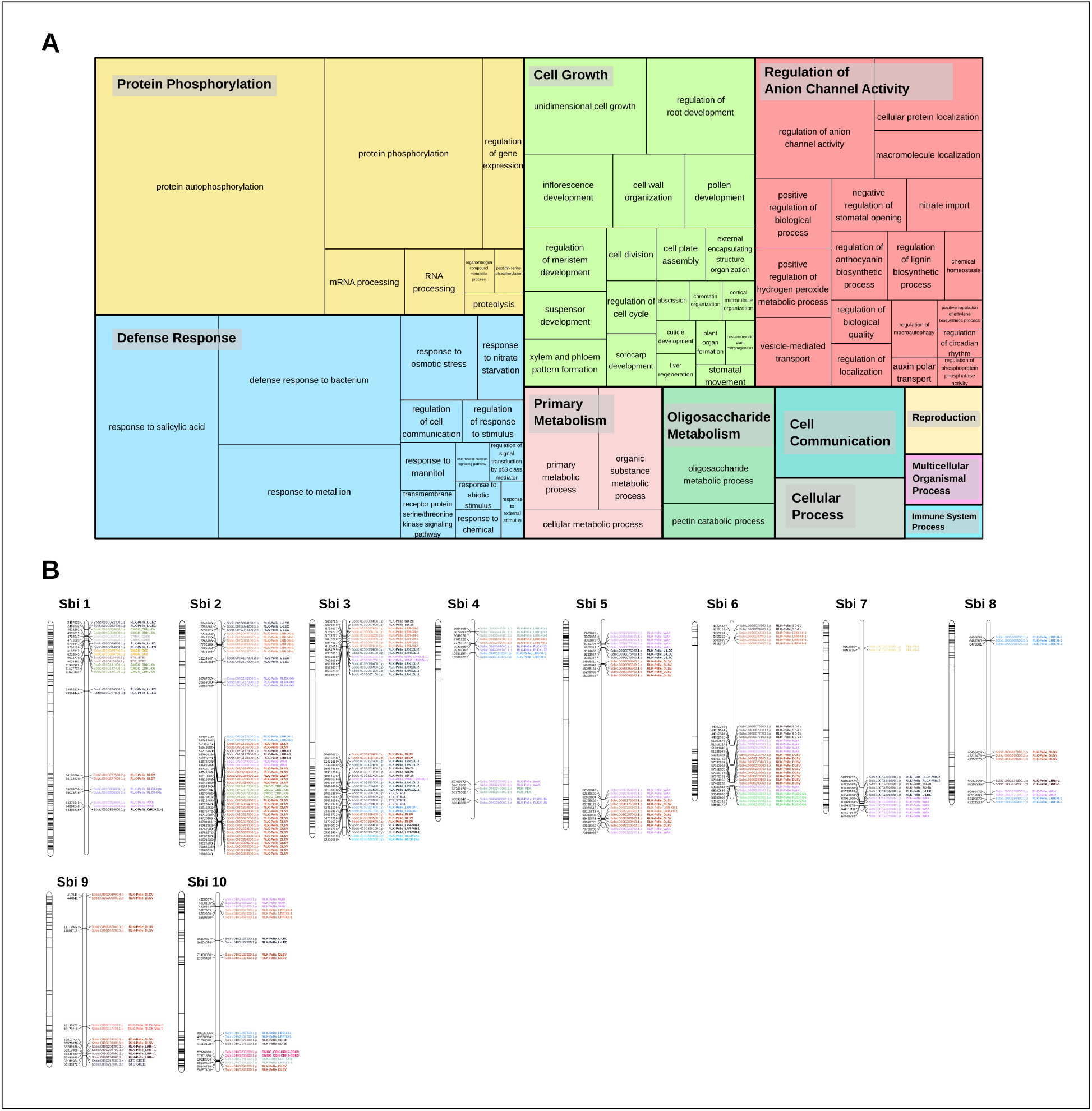
(A) Gene Ontology (GO) categories (biological processes) related to tandemly duplicated kinases in *Sorghum bicolor* (Sbi). The size of the subdivisions within the blocks represents the abundance of that category in this set of kinases. The colors are related to the similarity to a representative GO annotation for the group. (B) Kinase distribution along Ssp chromosomes. For each chromosome, all genes with kinase domains are indicated on the left, and only the tandemly organized kinases are indicated on the right, colored and labeled according to the subfamily classification.

In the Ssp genome, the distribution of PKs did not exhibit a clear pattern along chromosomes (Fig. 3B); however, in the Sbi genome, PK genes were concentrated in subtelomeric regions and were almost nonexistent in pericentromeric regions (Figure 4B). This pattern of distribution was observed more clearly when the tandemly distributed Sbi PKs were considered. Due to the importance of genes duplicated in tandem in biological processes, we also performed GO analysis to determine the categories related to tandemly duplicated kinases. The GO terms describing the biological processes of these proteins were clearly similar between Ssp and Sbi (Figs. 3A and 4A), and considerable correspondence to the total number of GO terms related to the entire set of PKs was observed (Supplementary Fig. S5A and B).

Segmental duplications accounted for the highest percentage of identified duplication types in both Sbi and Ssp PKs. The highest quantities in Ssp were observed in the allelic copies of chromosomes 1 and 2, which also contained the most PKs. In Sbi, chromosome 1 exhibited the most segmental duplications, even though chromosome 3 had the most PK genes. For all gene pairs within these collinear duplications, we calculated the Ka and Ks values to obtain a time indicator of these events and evaluated the primary influence of PK expansion by calculating the Ka/Ks ratio. We considered each gene pair to be under neutral (Ka/Ks=1), negative (Ka/Ks*<*1) or positive selection (Ka/Ks*>*1) (Zhang *et al*., 2006). The distribution of Ks values is visualized in Fig. 2E and F, and a full contrast is provided in Supplementary Tables S28 and S29. The Ks values were clearly more evenly distributed in Sbi than in Ssp, which had 1,287 (27.5%) segmentally duplicated PKs with a Ks of < 0.05. We used the Ks values to estimate the occurrence times of these duplications; the times ranged between 0 and 230.1 million years ago (MYA) in Ssp, with an average of 45.6 MYA, and between 4.9 and 229.7 MYA in Sbi, with an average of 96.8 MYA. Most segmental duplications with Ks<0.05 in Ssp were estimated to have occurred less than 3.83 MYA. Regarding the Ka/Ks ratio, we found the largest percentage of gene pairs as likely to be under negative selection in both species (∼86% in Ssp and ∼88% in Sbi).

All collinear duplications are shown in Fig. 5. The segmental events among alleles had different configurations, but in most duplications, the order of PKs on one allele was retained on the other allele (Fig. 5A). The correspondences among different chromosomes were much higher in Ssp (Fig. 5B) than in Sbi (Fig. 5C), mainly because of the allele specificity of Ssp, which is not known for Sbi. The duplication patterns were similar between Ssp and Sbi, and this genomic organization is clearly shown in Fig. 5D, where the kinase genomic correspondences indicate the increased synteny between these two species. In most PK subfamilies, the origin of most PKs was characterized by segmental duplications (109 subfamilies in Sbi and 115 in Ssp; Supplementary Tables S22 and S23). Interestingly, 4 subfamilies in Ssp (RLK-Pelle_RKF3, CMGC_Pl-Tthe, SCY1_SCYL2, and CMGC_GSKL) and 8 in Sbi (RLK-Pelle_RLCK-Os, PEK_GCN2, RLK-Pelle_RLCK-XI, STE_STE20-Pl, TLK, SCY1_SCYL2, TKL-Pl-8, and TKL-Pl-7) did not contain any PKs possibly originated by segmental duplications.

**Fig. 5:**
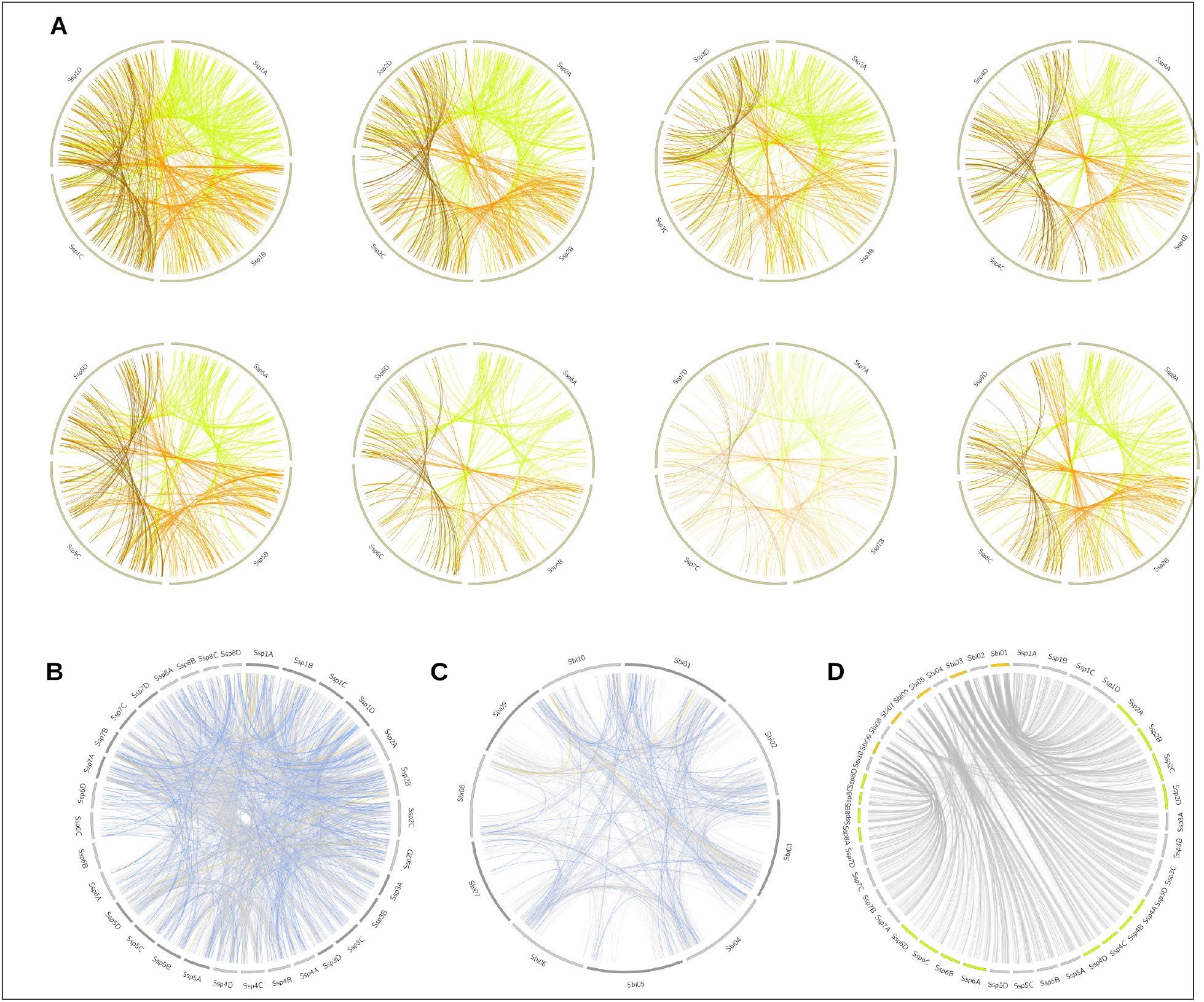
Segmental duplication events in the *Saccharum spontaneum* (Ssp) and *Sorghum bicolor* (Sbi) genomes, divided into (A) Ssp duplications between alleles on the same chromosome, with the colors representing the origin of the duplication (green for allele A, orange for allele B, and brown for allele C); (B) Ssp duplications between chromosomes, excluding events between alleles on the same chromosome; and (C) Sbi duplications. The colors in (B) and (C) indicate the selection type of the gene pair duplication (gray indicates negative selection; light blue, positive selection; and blue, neutral selection). (D) Representation of kinase correspondences between Sbi and Ssp, indicating the synteny relationships among these species.

Due to the PK allele specificity in Ssp, we performed additional analysis to assess the distribution of kinase copies among alleles and investigated possible associations among allelic copies, duplications and related domains (Fig. 6). Each Ssp GM can have up to four allelic copies, depending on the genomic organization of the gene. Subfamilies with larger numbers of PKs had a more dispersed organizational profile in terms of the number of allelic copies per GM. Subfamilies with fewer GMs, on the other hand, did not have a uniform configuration. These subfamilies constitute the majority of the Ssp kinome (∼60% of the subfamilies had 5 or fewer representative GMs, and 33 subfamilies (∼30%) had only 1 GM). Even with the few related proteins, these small subfamilies did not exhibit similar characteristics. Only 3 of these GMs had copies on the 4 alleles, 10 GMs contained copies on 3 alleles, 9 on 2 alleles, and 11 on only one allele (3 in allelic model A, 3 in B, 2 in C and 3 in D). More tandem and segmental duplications were clearly observed in subfamilies with more elements, but this pattern did not hold for the quantity of functional domains and multikinase domains. Even though the subfamilies foremost clearly exhibiting these characteristics have already been described, these results are further supported in Fig. 6.

**Fig. 6:**
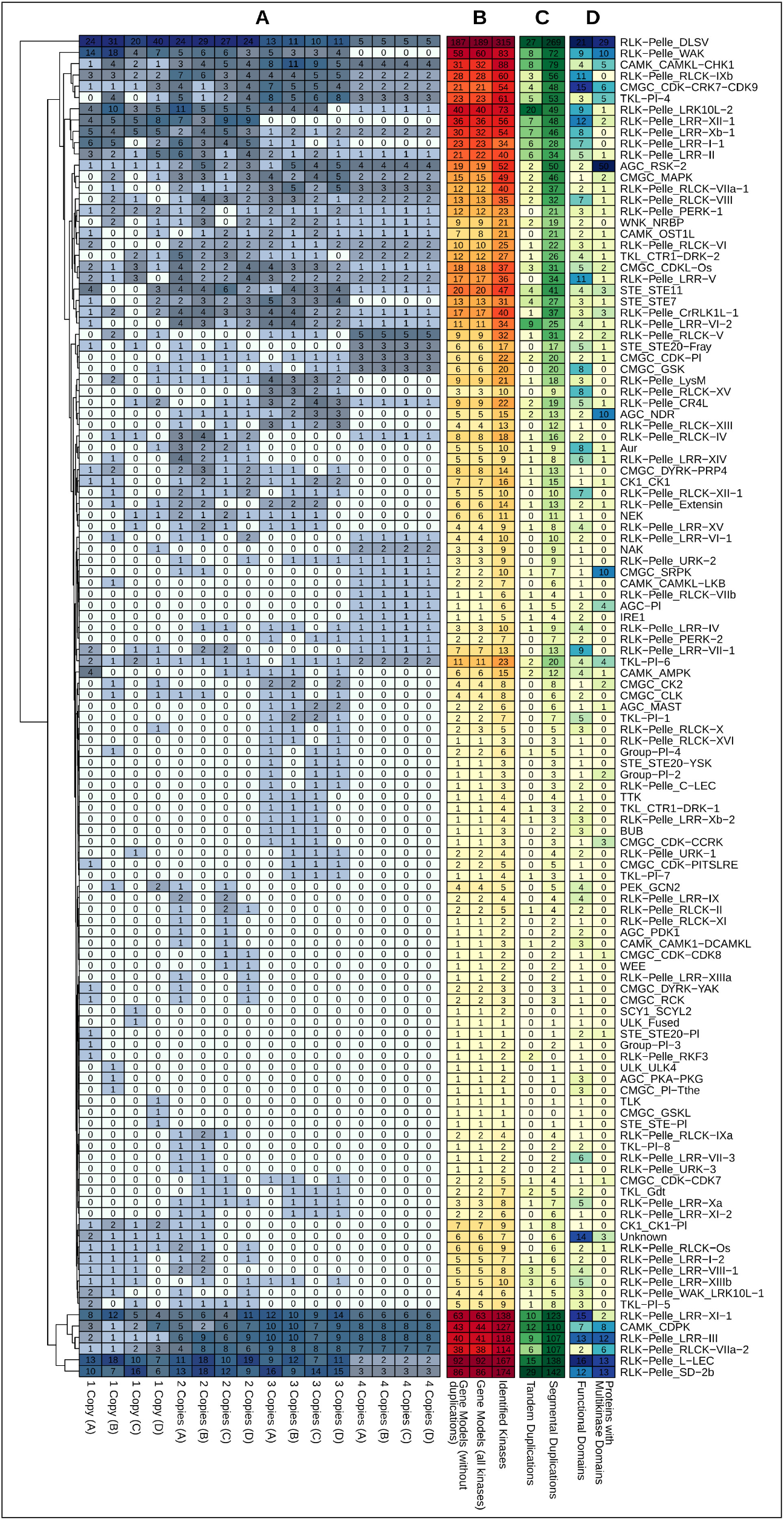
Heatmap representations of kinase subfamily profiles in *Saccharum spontaneum* related to (A) kinase copies among alleles; (B) subfamily quantification considering the entire set of kinases and the respective quantity of gene models; (C) tandem and segmental duplication events; and (D) the presence of different functional domains and multikinase domain-containing proteins within subfamilies.

### 3.4. Estimates of kinase expression and construction of coexpression networks

Quantification of kinase expression in Sbi and Ssp was performed via a wide variety of datasets and comprised different tissues and genotypes (Supplementary Tables S1 and S2). For each species, the bioinformatic procedures included quality filtering of the raw sequencing reads followed by transcript quantification using Salmon software with the total set of Sbi and Ssp CDSs. From the CDS quantifications, we separated the subset of kinase coding genes. Via TPM values, Sbi kinase expression was quantified in 205 samples (Supplementary Table S30); Ssp kinase expression, in 234 (Supplementary Table S31). To quantify expression at the subfamily level, the TPM values for all PK members in a subfamily were averaged in each sample (Supplementary Tables S32 and S33). However, most of the experiments contained several biological and technical replicates, and the sample TPM values were also averaged to separately represent the unique characteristics of a tissue from a specific genotype (Supplementary Tables S34 and S35).

The expression quantification of Ssp and Sbi kinase subfamilies was visualized with a heatmap (Fig. 7). Evident distinctions are visible in the heatmap columns. Considering the hierarchical clustering analysis performed combining genotypes and tissues (columns) from both species, there was a noticeable division into 5 groups, also identified by the total within sum of squares using a range of group configurations (2-10). From right to left in the heatmap, the groups are separated into (I) sugarcane samples from internodes and roots; (II) Sbi samples from internodes, roots and spikelets; (III) Sbi samples from epidermal tissues, seeds and microspores; (IV) Sbi and sugarcane samples from leaves and shoots; and (V) Sbi samples from pollen. The expression patterns of kinase subfamilies were more similar among similar tissues from different species than among different tissues from the same species. However, these clusters contained subdivisions supporting the species specificities.

**Fig. 7:**
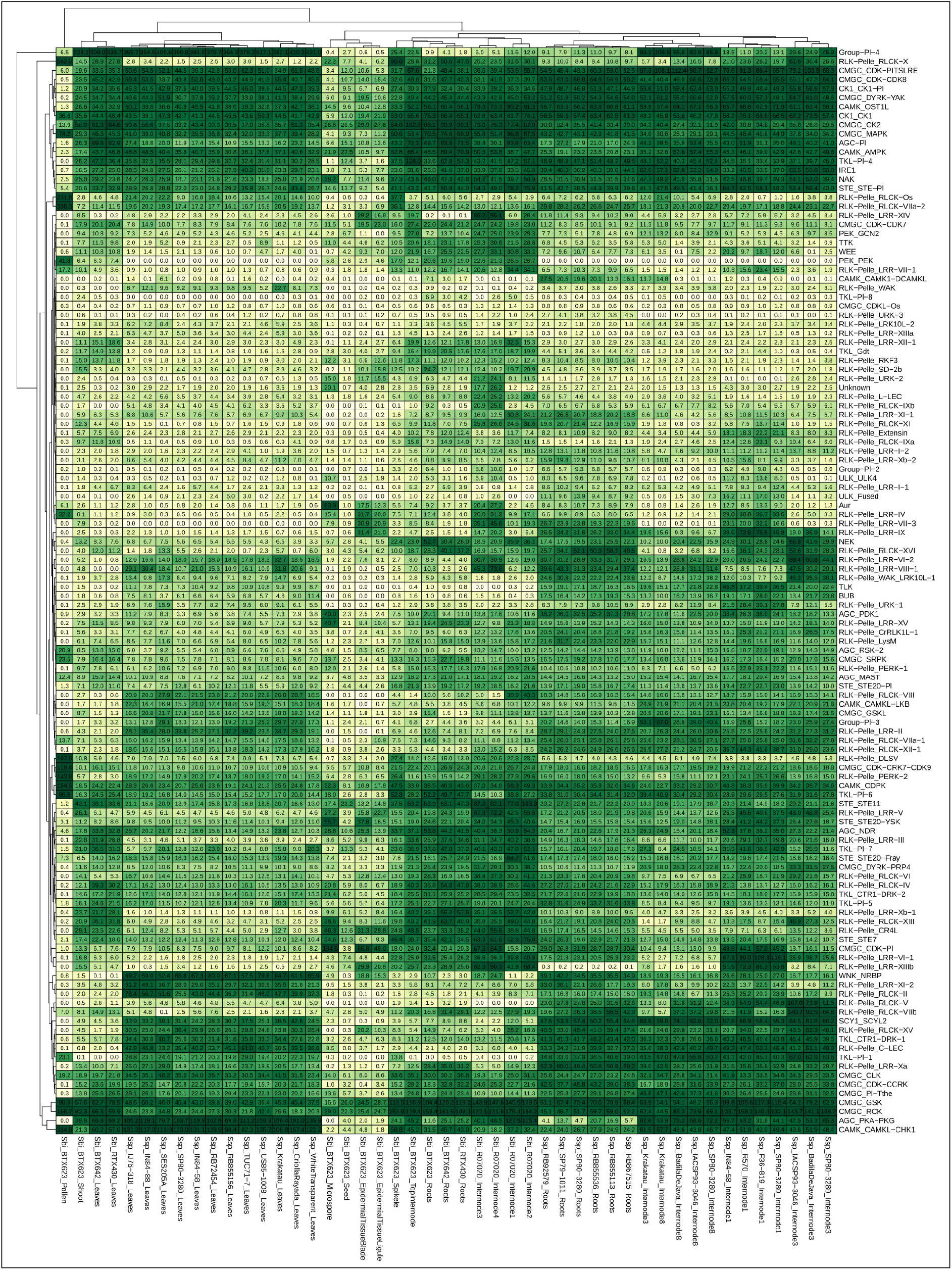
RNA expression profiles of *Saccharum spontaneum* and *Sorghum bicolor*, shown on a heatmap indicating the average sample values of different combinations of genotypes and tissues (columns) and considering the organization of kinase subfamilies (rows).

The differences in subfamily expression profiles were investigated further. For each family, we calculated the dispersion of expression among genotypes and tissues by using the statistical measures of the standard deviation and coefficient of variation (Supplementary Tables S36 and S37). The divergence of these measures among tissues was high in Sbi, as observed in the heatmap and indicated by the high values of the coefficient of variation (ranging from ∼38% to ∼297%). In Ssp, on the other hand, 16 subfamilies exhibited relatively uniform expression patterns in the analyzed samples (with coefficient of variation of less than or equal to 20%). The coefficients of the other subfamilies ranged from ∼21% to ∼213%. This difference is possibly explained by greater diversity of tissues used for Sbi than for Ssp. To identify the kinase subfamilies with the highest and the lowest expression values, we calculated additional statistical measures to summarize the distribution of TPM values in each subfamily (i.e., minimum, maximum, mean, and 1st, 2nd and 3rd quartiles). We selected the 12 subfamilies (10% of the dataset) with the highest and lowest values of all statistical measures employed. We considered a subfamily as having the highest or the lowest expression values if that subfamily was ranked in at least 4 of the 6 measures. We identified 8 subfamilies in Sbi and 12 in Ssp with the highest expression patterns in the dataset. Surprisingly, three of these subfamilies (CK1_CK1, CMGC_GSK, and CMGC_RCK) had the highest expression value in both species. Even though their expression values were significantly increased, these subfamilies did not contain the highest numbers of kinases (CMGC_RCK, for example, contained only 3 members in both Sbi and Ssp). In Sbi, we also found CAMK_CDPK, CMGC_CK2, CMGC_MAPK, RLK-Pelle_RLCK-X, and STE_STE11; and in Ssp, AGC_PKA-PKG, CAMK_CAMKL-CHK1, CAMK_OST1L, CK1_CK1-Pl, CMGC_CDK-CDK8, CMGC_CDK-PITSLRE, CMGC_DYRK-YAK, Group-Pl-4, and SCY1_SCYL2.

By this approach, 8 subfamilies in Ssp and 9 in Sbi with the lowest expression values were identified, with two overlapping subfamilies (CMGC_CDKL-Os and RLK-Pelle_URK-3). In Sbi, we also identified BUB, CAMK_CAMK1-DCAMKL, RLK-Pelle_LRR-I-1, RLK-Pelle_RLCK-V, RLK-Pelle_WAK, TLK, and ULK_Fused. In Ssp, we found Group-Pl-2, RLK-Pelle_LRR-XIIIa, RLK-Pelle_URK-2, TKL_Gdt, TKL-Pl-8, and ULK_ULK4. RLK-Pelle_URK-3 had only one kinase member in both the Sbi and Ssp kinomes; however, CMGC_CDKL-Os had 37 kinases in Ssp and 23 in Sbi. Due to the apparent lack of a correlation between the expression values and the numbers of kinases in the subfamilies, we calculated the Spearman correlation coefficient between the subfamily expression estimates and kinase quantities (Supplementary Tables S38 and S39), and we did not find any combination of genotype/tissue with a significant correlation—even when only genes with tandem or segmental duplications were compared.

Together, the dendrogram and the heatmap indicate the presence of groups of subfamilies with high similarities, whose expression patterns changed jointly according to the tissue/genotype. Collectively considering all Sbi and Ssp quantifications, we evaluated their similarities through correlation analysis. The strongest correlations were higher than 0.97 for the two subfamily pairs RLK-Pelle_RLCK-Os/RLK-Pelle_RLCK-VIIa-2, and RLK-Pelle_RLCK-VIIa-2/RLK-Pelle_RLCK-X. However, to expand and complement the assessment of the similarities in RNA expression among the subfamilies, we also constructed coexpression networks based on the expression correlation among samples in each subfamily. Two networks were constructed: one for Sbi and one for Ssp (Fig. 8). Each node in the network represents a different kinase subfamily (the node sizes represent the mean of the expression values within the subfamily) and each connection has a minimum Pearson correlation coefficient of 0.6 (the edge sizes represent the degree of the correlation). With the network structure, we evaluated the presence of cohesive clusters formed by correlated subfamilies using a network community detection approach based on label propagation. In the Sbi network (Fig. 8A), we identified 4 different modules with 87, 15, 3 and 9 elements. Four modules were also identified in the Ssp network (Fig. 8C), but the distribution of the elements differed (83, 13, 8, and 2). In both networks, some subfamilies (6 in the Sbi network and 13 in the Ssp network) were identified as disconnected elements without any significant relationship with the other elements of the networks. There was no evident similarity between these communities (Supplementary Fig. S6, Supplementary Tables S40 and S41), indicating the differences in the expression pattern correspondences between Ssp and Sbi.

**Fig. 8:**
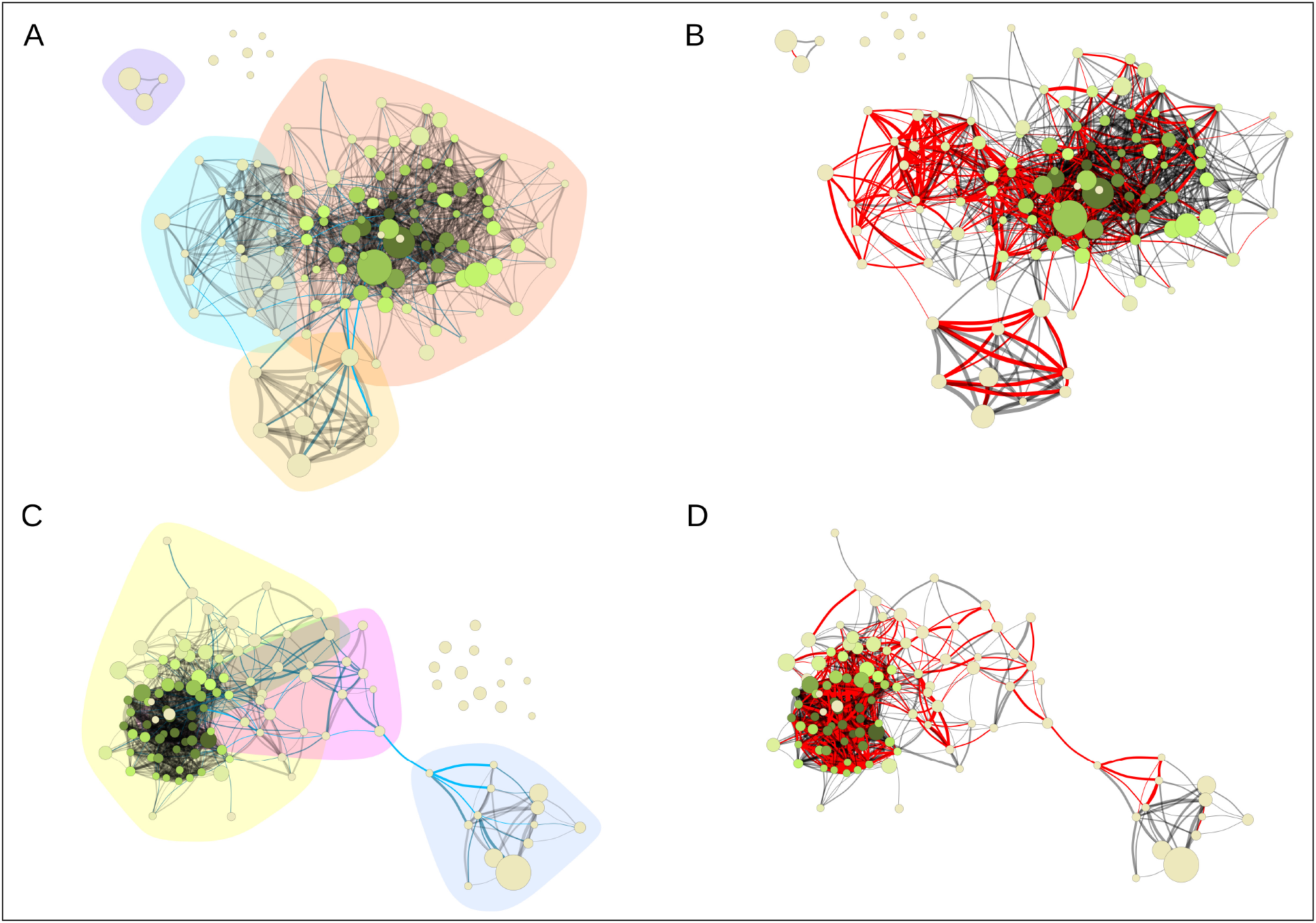
Coexpression networks for *Sorghum bicolor* (Sbi) and *Saccharum spontaneum* (Ssp) kinase subfamilies. Each node corresponds to a different subfamily, its size corresponds to the average expression value for all kinases within the subfamily in different samples, and its color corresponds to the hub score and ranges from beige to dark green. Each edge corresponds to a correlation with a Pearson correlation coefficient of at least 0.6. The correlation strength is represented by the edge’s width and the edge betweenness score is represented by the color (ranging from black to light blue, with light blue representing the highest values). (A) Sbi network with the background colored according to the community detection analysis. (B) Sbi network indicating the similarities with the Ssp network in red. (C) Ssp network with community structure information. (D) Ssp network indicating the similarities with the Sbi network in red.

Apparently, the Sbi and Ssp networks exhibited many different forms and structures; however, by highlighting the connections in common between the Sbi and Ssp networks (Fig. 8B and D), we observed a similar and important substructure between these representations. The main network components were connected by this core structure, indicating the strongest correlations between kinase subfamilies. The Ssp network (Fig. 8D) contained an edge that clearly separates the network into two components; interestingly, this edge also belonged to the common structure. By coloring the network edges according to the betweenness measure (Fig. 8A and B), we defined the connections between subfamilies that were most likely to represent vulnerabilities in the networks, possibly indicating influential subfamilies in this complex system. The most important connections were related to the subfamily pairs CAMK_CDPK/RLK-Pelle_LRR-VI-2 and CAMK_CDPK/CMGC_RCK in Sbi, and to RLK-Pelle_L-LEC/RLK-Pelle_LRR-VIII-1, RLK-Pelle_CR4L/RLK-Pelle_LRR-VIII-1, and RLK-Pelle_CR4L/RLK-Pelle_LRR-Xb-1 in Ssp (Supplementary Tables S42 and S43).

The last analysis we performed on the networks aimed to identify the most influential subfamilies by ranking the nodes according to their hub scores. These scores were used to color the nodes in the network (the correspondences between nodes and subfamilies are indicated in Supplementary Figs. S7 and S8 and Supplementary Tables S40 and S41). The highest hub scores denote kinase subfamilies with the most connections in the network. The top 5 scores belonged to the subfamilies RLK-Pelle_LRR-III, RLK-Pelle_RLCK-XII-1, CMGC_CDK-CRK7-CDK9, CMGC_GSK, and RLK-Pelle_Extensin in Ssp, and to the Unknown category, RLK-Pelle_LRR-XV, CMGC_GSK, STE_STE20-Fray, and CAMK_OST1L in Sbi. Additionally, high expression values in subfamilies did not indicate increased hub scores (Fig. 8).

## 4. Discussion

Sugarcane possesses one of the most complex genomes known among crops (O’Hara and Mundree, 2016; Mancini *et al*., 2018; De Souza Barbosa *et al*., 2020) which could, until recently, only be assembled at the scaffold level (Grativol *et al*., 2014; Okura *et al*., 2016; Riaño-Pachón and Mattiello, 2017; Souza *et al*., 2019). Only in 2018 did an approach coupling next-generation and long-read sequencing with high-throughput chromatin conformation capture enable a chromosome- and allele-level genome of a Ssp clone to be assembled (Zhang *et al*., 2018). This study paved the way for several comprehensive analyses of gene families in this species (Hu *et al*., 2018; Hua-Ying *et al*., 2019; Lin *et al*., 2019; Shi *et al*., 2019; Wang *et al*., 2019a, b, c; Zhang *et al*., 2019b; Feng *et al*., 2020; Huang *et al*., 2020; Li *et al*., 2020b; Su et al., 2020a, Zhang and Yin, 2020). In the present work, however, we analyzed the kinome of not only sugarcane but also sorghum, a close diploid relative. Studies estimate that the *Saccharum* and *Sorghum* lineages diverged 4.6-5.4 MYA (Kim *et al*., 2014). After diverging from *Miscanthus* 3.1-4.6 MYA (Kim *et al*., 2014), the *Saccharum* lineage experienced at least two rounds of whole-genome duplication (Zhang *et al*., 2018), while Sbi remained diploid. Therefore, sorghum genomic resources are a valuable resource in genetic studies in sugarcane (Paterson *et al*., 2009), in which they have been extensively employed (Okura *et al*., 2016; Mancini *et al*., 2018; Bedre *et al*., 2019). As the genomes of both species are now available, comparisons of the diversity, organization and expression of PKs between these two species enable us to perform in-depth explorations of the evolutionary history of these proteins, which are relevant to numerous biological processes.

A previous work that classified PKs from 25 plant species (Lehti-Shiu and Shiu, 2012) indicated that substantial numerical variations in this superfamily exist among species; however, this variation could be overestimated due to differences in the completeness of the genomic assemblies. Moreover, the estimates presented by this and other studies (Singh *et al*., 2014; Wei *et al*., 2014; Liu *et al*., 2015; Zhu *et al*., 2018a, b) indicated a number of PKs in Sbi (1,210) very similar to those of other Poaceae species, which range between 1,041 in Bdi to 1,417 in Osa (Lehti-Shiu and Shiu, 2012; Wei *et al*., 2014). This figure is also comparable to the 1,093 PK genes previously estimated for this species using an earlier genomic reference (Lehti-Shiu and Shiu, 2012). The Ssp genome, on the other hand, contains one of the largest numbers of PK genes reported for any plant species (2,919), ranking below only the allohexaploid genome of *Triticum aestivum* (3,269 PKs) (Yan *et al*., 2017). However, we must consider that this identification was performed using a genome that provides information at the allele level; when only Ssp GMs (i.e., single representatives of all copies of a gene) were analyzed, we found a much lower number of PK genes (1,345), which is also within the range of PKs in other Poaceae species. This discrepancy reinforces the hypothesis of Lehti-Shiu and Shiu (2012) that the expansion of PK genes is directly related to recent whole-genome duplication events, a suggestion that was made considering that paleopolyploid species, such as soybean, have larger repertoires of these proteins. Indeed, because soybean’s duplication events are much more ancient than sugarcane’s (having occurred ∼13-59 MYA) (Schmutz *et al*., 2010), its homologous chromosomes are not treated as allelic copies. Therefore, it is only natural that more PK genes were identified in the two kinomes compiled for Sbi, namely, 2,099 (Lehti-Shiu and Shiu, 2012) and 2,166 (Liu *et al*., 2015) PKs, while Ssp, which underwent very recent polyploid evolution, contained many fewer PK genes when allelic copies were considered.

In Sbi, PK genes were more commonly located in subtelomeric regions. This pattern was even more evident when only tandemly duplicated PKs were considered; similar (though less pronounced) patterns were observed in the kinomes of soybean (Liu *et al*., 2015), *T. aestivum* (Yan *et al*., 2017), *Gossypium raimondii* and *Gossypium barbadense* (Yan *et al*., 2018). Yan *et al*. (2017) noted that this pattern is consistent with *T. aestivum*’s higher gene and expressed sequence tag (EST) densities in distal regions of chromosomes and inferred that this location pattern of PKs could indicate chromosomal rearrangements. Our findings are equally compatible with the general genomic landscape of sorghum: in this species, the density of genes—especially paralogs—is much higher in chromosome extremities, while pericentromeric regions are very rich in long terminal repeat retrotransposons (Paterson *et al*., 2009; Mace and Jordan, 2011). Studies have further demonstrated that genes are not uniformly distributed throughout the Sbi genome but rather clustered in regions termed “gene insulae” (Gottlieb *et al*., 2013).

On the other hand, the gene density in Ssp is less skewed towards subtelomeric regions (Zhang *et al*., 2018), which might explain why we did not observe such a clear pattern of PK gene distribution along Ssp chromosomes. An analogous observation was made in a recent analysis that compared the genomic structure of Sbi and those of two *Saccharum* species (Zhang *et al*., 2019a). Although the three species exhibited considerable collinearity among homologous chromosomes, genes that were widely dispersed in *S. officinarum* and *S. robustum* linkage groups were much more tightly clustered in subtelomeric regions on Sbi chromosomes. This same pattern is visible on circos plots that show the synteny between the Sbi and Ssp kinomes (Fig. 5D); although many of the Sbi PK genes are also present in the Ssp genome and are much more widely distributed along chromosomes in Ssp. The dispersion of Ssp kinase genes between chromosomes and allelic copies was also relatively balanced and somewhat proportional to the chromosome length. Overall, this is similar to the patterns of kinase genes reported for rice (Dardick *et al*., 2007); pineapple and grapevine, although these genes are more unevenly distributed along chromosomes (Zhu *et al*., 2018a, b).

Comparison of the Sbi and Ssp kinomes alone reveals that their subfamily composition profiles are very similar. The only subfamily found exclusively in Sbi was PEK_PEK. Even though the PEK family is responsible for phosphorylation of eukaryotic translation initiation factor 2 subunit alpha (eIF2*α*) (Immanuel *et al*., 2012), each subfamily is involved in the response to different types of stresses (Donnelly *et al*., 2013). The PEK_GCN2 subfamily was found in both species, and its activation is related to amino acid and glucose deprivation (Yang *et al*., 2000; Deval *et al*., 2009; Baker *et al*., 2012), viral infection (Berlanga *et al*., 2006; Krishnamoorthy *et al*., 2008) and UV irradiation (Grallert and Boye, 2007). PEK_PEK subfamily kinases are especially activated during endoplasmic reticulum (ER) stress (Baker *et al*., 2012), and are homologous to IRE1 subfamily proteins (Urano *et al*., 2000), which are also activated in response to ER stress (Liu *et al*., 2007a) and were found in both Sbi and Ssp kinomes.

The group containing the most kinases is the RLK-Pelle group (Gish and Clark, 2011) and, similar to findings in other kinomes (Singh *et al*., 2014; Wei *et al*., 2014; Zulawski *et al*., 2014; Liu *et al*., 2015; Yan *et al*., 2017, 2018; Zhu *et al*., 2018a, b), we found that the RLK-Pelle group had the most members in our study. This expansion in the Sbi and Ssp kinomes is apparently related to a few specific families within this group, most notably the LRR, RLCK, DLSV, L-LEC and SD-2b families. These families have already been associated with the increased number of kinases in the RLK-Pelle group (Zhu *et al*., 2018a, b), mostly because of their relation with biotic and abiotic stress responses (Dezhsetan, 2017). In the cotton kinome, for instance, LRR subfamilies have been suggested to be significantly associated with plant growth, development and defense responses (Yan *et al*., 2018), and these associations have already been described for Sbi and Ssp. In Sbi, the LRR family has broadly been linked with the response to several types of stress (Kawahigashi *et al*., 2011; Azzouz-Olden *et al*., 2020; Filiz and Kurt, 2020), playing roles related to signal transduction in response to extracellular signals (Azzouz-Olden *et al*., 2020; Dhaka *et al*., 2020; Vikal *et al*., 2020), pollen development (Dhaka *et al*., 2020), metabolism, and chaperone functions (Vikal *et al*., 2020). In Ssp, in addition to its association with defense response processes (Xu *et al*., 2018; Yang *et al*., 2019), this family has already been associated with hormone metabolism (Chen *et al*., 2019), cellulose and lignin biosynthesis (Kasirajan *et al*., 2018), and sucrose synthesis (Vicentini *et al*., 2009).

In addition to the remarkably important LRR family, the RLCK, DLSV, L-LEC, and SD-2b families are also involved in diverse essential mechanisms. Because RLCK family members do not contain extracellular and transmembrane domains (Gao and Xue, 2012; Zulawski *et al*., 2014), these proteins are generally involved in more specific processes (Jurca *et al*., 2008). In addition to disease resistance, RLCK proteins have been shown to be related to plant growth, immune responses (Yan *et al*., 2018; Zhu *et al*., 2018a), and vegetative development (Jurca *et al*., 2008; Gao and Xue, 2012). Together with RLCK family members, DLSV family members were found to be differentially expressed in soybean tissues in stress experiments (Liu *et al*., 2015). The DLSV family includes Domain of Unknown Function 26 (DUF26), SD-1, LRR-VIII, and VMA (a RLK subfamily specific in moss)-like proteins (Lehti-Shiu and Shiu, 2012), which mediate the control of stress responses and development (Vinagre *et al*., 2006; Vaattovaara *et al*., 2019), with some members being associated with signaling pathways regulating the responses to cold (Yan *et al*., 2017) and infection (Yan *et al*., 2017). L-LEC and SD-2b have established associations with the defense response (Chen *et al*., 2006; Wei *et al*., 2014) but also with stomatal immunity regulation via an L-LEC member (Desclos-Theveniau *et al*., 2012) and with self-incompatibility via SD-2b (Stein *et al*., 1991). The essentiality of mechanisms shared by these families clearly indicates their functional importance among plants (Vaattovaara *et al*., 2019) and demonstrates their importance in the expansion and maintenance of the Sbi and Ssp kinomes.

Differences in PK composition may lead to different functional profiles. Similarly, structural divergences may arise at distinct points in evolutionary history (Teich *et al*., 2007; Liu *et al*., 2015), contributing to different domain organizations and, subsequently, to diverse functions (Xu *et al*., 2012). Although PKs in the same subfamily have similar intron distribution profiles in wheat (Yan *et al*., 2017), several compositional differences were detected in the soybean kinome (Liu *et al*., 2015); we also detected these differences in the Ssp and Sbi kinomes. In the Sbi kinome, the distribution of introns across subfamilies was more organized than that in the Ssp kinome, indicating the more recent intron/exon reorganization of Ssp PKs. This more evidently cohesive structure among Sbi subfamilies than Ssp subfamilies indicates that gene reorganization may have occurred after these species diverged. The NEK, CK1_CK1-Pl, PEK_GCN2, and TKL_CTR1-DRK-2 families had the most prominent structural organization in both Sbi and Ssp. All of these families play essential roles in cellular processes, which requires a higher level of organization. As mentioned previously, PEK_GCN2 activity is linked with eIF2*α*. Interestingly, the Ssp kinome contained more members of this family than any other species examined (Supplementary Fig. S4), and this family had a considerable gene organization. NEK family members have been associated with the cell cycle machinery through microtubule organization, cell growth, and stress responses (Moniz *et al*., 2011; Takatani *et al*., 2015). The CK1_CK1-Pl subfamily is part of the CK1 group identified by Pei *et al*. (2019), which is involved in several vital physiological processes via phosphorylation of different substrates (Tan and Xue, 2014; Karpov *et al*., 2019). TKL_CTR1 members have already been linked to defense response pathways, including the ethylene signal transduction pathway (Varberg *et al*., 2018).

In contrast with the highly organized gene profile found in some subfamilies, several sets of PKs exhibited considerable domain diversity and composition. Ssp PK subfamilies had the largest number of domains, corroborating the most recent possible gene organization of these PKs. RLK-Pelle subfamilies showed the largest differences in domain composition in both Sbi and Ssp, as expected due to the large size of this family. In addition to RLK members, the CMGC_CDK-CRK7-CDK9 (in Ssp) and CMGC_GSK subfamilies were among the top 10% of subfamilies with the largest number of different domains. Even though the CMGC_CDK-CRK7-CDK9 subfamily had the most members among the CMGC group, the number of PKs in the CMGC_GSK subfamily was similar to those in the other GMC subfamilies. Therefore, this domain diversity might be explained by the diverse functions performed by these proteins. As previously mentioned, the RLK-Pelle group putatively participates in a wide variety of induced biological processes, and the CMGC_CDK family (Joubès *et al*., 2000) also integrates several functions of transcription and cell division (Malumbres, 2014). Specifically, the CRK7 and CDK9 subfamilies are related to the numerous processes in cell cycle control (Goldberg *et al*., 2006). Additionally, the GSK subfamily affects numerous signaling pathways (Wrzaczek *et al*., 2007).

Another interesting observation relates to the number of potential PK genes that were not considered because they had a domain coverage of less than 50%, indicating that they represent atypical kinases or pseudogenes (Lehti-Shiu and Shiu, 2012; Liu *et al*., 2015). For Sbi, this criterion resulted in the exclusion of 57 genes, which accounted for ∼3% of sequences with significant correspondences with PKs. In Ssp, however, 735 such genes were discarded, accounting for almost 20% of the initially identified PKs. In their pioneering work, Lehti-Shiu and Shiu (2012) found that 9.6% of all kinases initially identified in 25 species exhibited a domain coverage of less than 50%, and this value varied considerably in later studies that employed the same methodology. We can also speculate on the influence of polyploidization on the pseudogenization of PK genes. While no kinomes have been published for other autopolyploid species, we may take as examples those that have been generated in allopolyploids. In the *Triticum*-*Aegilops* complex, the kinome of the allohexaploid *T. aestivum* contains ∼22% atypical kinases, while the kinomes of two of its diploid parental species, *Triticum urartu* and *Aegilops tauschii*, contain ∼16 and ∼14% atypical kinases, respectively. Similarly, in *Gossypium*, the kinomes of two diploid species (*G. raimondii* and *Gossypium arboretum*) contain ∼4 and ∼9% atypical kinases, while in the kinomes of the allotetraploids *Gossypium hirsutum* and *G. barbadense*, ∼12% of PKs have these characteristics. The larger numbers of kinase genes with atypical domains in polyploid genomes may have resulted from more frequent pseudogenization events in these species and subsequent whole-genome duplication (WGD), a long-proposed consequence of gene duplication and thus of polyploidization (Magadum *et al*., 2013).

Even though multikinase domains were found in both Sbi and Ssp, Ssp contained more than Sbi, and more repetitions were found in some PKs. Similar to the soybean and grapevine kinomes (Liu *et al*., 2015; Zhu *et al*., 2018b), the Sbi kinome contained PKs with only 2 or 3 kinase domains, in contrast to the Ssp kinome, which contained PKs with between 2 and 5 kinase domains. Interestingly, the AGC_RSK-2 subfamily was found to have the largest number of multikinase domains in both Sbi and Ssp, accounting for a very high percentage of members of this subfamily, which is explained by the fusion of two PKs in the evolutionary history of the RSK family (Carriere *et al*., 2008). The AGC_NDR subfamily also exhibited this notable characteristic; however, in this subfamily, the large number of multikinase domains is associated with the insertion of a nuclear localization signal within the kinase domain (Tamaskovic *et al*., 2003). Moreover, in the Sbi and Ssp kinomes, the PKs with most kinase domains were in the RLK-Pelle_WAK subfamily, which is functionally linked to cell growth (Gish and Clark, 2011) and whose loss might result in lethality (Wagner and Kohorn, 2001). As the percentage of multikinase domains found in this family was small, we consider that such domains may interact with specific substrates (Liu *et al*., 2015).

Our study is the first to categorize a kinase superfamily considering allele copies. Even though the presence of kinase domains in Ssp PKs was highly conserved, differences in intron exon organization and domain composition were found. The most common compositional differences were related to domain distribution along the allele copies (e.g., inversion of LRR and kinase domains along the sequences), insertion or loss of domains in allele copies (e.g., LRR, antifungal, and uroporphyrinogen decarboxylase domains, as well as domains of unknown function); and duplication of domains (e.g., LRR, legume lectin, EF-hand, and kinase domains). Even though we expected minor differences across allele copies, these findings suggest specific rearrangements of kinase sequences, indicating possible functional associations. Other studied protein families in Ssp also had different pattern distributions across allele copies (Huang *et al*., 2020; Li *et al*., 2020b). In some studies, the gene structure has been reported to be similar across these copies; however, this pattern is not universal (Ma *et al*., 2019; Shi *et al*., 2019). The genomic structure and organization of sugarcane is considerably complex (Sforça *et al*., 2019), and the pattern of gene distribution across alleles is unclear; thus, more studies on specific genes and subfamilies are required to better understand the organization of the sugarcane genome.

We also performed several *in silico* analyses to evaluate the molecular characteristics of the PKs identified in the two species. As reported for grapevine (Zhu *et al*., 2018b), the pIs and MWs of the PKs were generally similar within subfamilies in Ssp and Sbi; these results were expected, as these properties are estimated based on the protein sequence. We observed, however, that Ssp contained many more PK subfamilies with significant variation in the MW than did Sbi, possibly indicating a broader diversity of kinases in Ssp. After verifying the presence of signal peptides in the PK sequences, we estimated that ∼40% of Sbi kinases contained signal peptides, in contrast with ∼30% in Ssp. This percentage is very similar to that in maize, where ∼30% of PKs contain these signal sequences (Wei *et al*., 2014). Regarding the subcellular localization of the PKs, we noted high divergence in the results obtained with the tested tools. All three methods (Yu *et al*., 2006; Horton *et al*., 2007; Sperschneider *et al*., 2017) are based on machine learning techniques and have unique advantages. Therefore, the discordant localizations may not be reliable, and we decided to use a consensus approach, considering only the results consistent between at least two of the tools. Although this process did not allow the subcellular localization of all PKs to be estimated, it did allow us to determine a more consistent predictive set for categorizing the Sbi and Ssp kinomes. Due to this conservative approach, we did not make inferences about the distribution of subfamily localizations.

Annotation of the PKs based on GO terms corroborated the accuracy of their identification. For instance, in both the Ssp and Sbi kinomes, the five most frequently appearing annotated GO terms were (I) defense response to oomycetes, (II) protein serine/threonine phosphorylation, (III) binding, (IV) plasma membrane and (V) pollen development (Fig. 2C and D). All of these terms can be easily linked to kinases; indeed, terms (II) and (III) exhibit the most obvious associations, as PKs catalyze the phosphorylation of proteins by transferring terminal phosphate groups from ATP to serine, threonine or tyrosine residues in other proteins—a process that involves the binding of PKs to their targets (Hunter, 1995). A large portion of eukaryotic plant kinases (as we further demonstrated in the present work) are grouped into the RLK superfamily and are thus located in the plasma membrane, which explains term (IV). Additionally, PKs are frequently shown to participate in responses to infection by various oomycetes in many plant species (Hall *et al*., 2007; Blanco *et al*., 2008; Hok *et al*., 2011, 2014; Carella *et al*., 2019), as well as in pollen development, in several plants (Zhang *et al*., 2001; Xu *et al*., 2011; Lafleur *et al*., 2015; Chen *et al*., 2016; Li *et al*., 2018a), explaining terms (I) and (V). This logic was maintained when the annotation results were summarized in treemaps (Supplementary Fig. S5A and B); first, terms associated with protein phosphorylation were strongly represented in the kinomes of both species. This summarization also highlights the broad presence of terms associated with other mechanisms in which plant PKs are widely and historically known to be involved, such as defense responses (Chen *et al*., 2006; Tena *et al*., 2011; Wei *et al*., 2014; Xu *et al*., 2018; Yang *et al*., 2019), cellular development (Jin *et al*., 2002; Matschi *et al*., 2013; Komis *et al*., 2018), regulation of stomatal closure (Li *et al*., 2000; Mustilli *et al*., 2002; Lee *et al*., 2016) and development of leaves and pollen (Roe *et al*., 1993; Benjamins *et al*., 2001; Khew *et al*., 2015).

In Sbi, we also investigated 100 PKs that are possibly subject to alternative splicing, a process that leads to the production of different mRNA isoforms from a single gene, therefore expanding the functional diversity of the gene. Alternative splicing is extensively reported to regulate plant development, circadian clocks and responses to environmental stimuli, especially stresses (Filichkin *et al*., 2015; Shang *et al*., 2017). When only alternatively spliced PKs were annotated and summarized (Supplementary Fig. 5C), we observed similarities to the categories associated with all GO terms in the two species analyzed. One notable difference was the inclusion of a category that included terms related to programmed cell death, a stress-triggered process (Danon *et al*., 2000) controlled by PKs (Tang *et al*., 2005; Liu *et al*., 2007b; Lachaud *et al*., 2013; Wrzaczek *et al*., 2014; Yadeta *et al*., 2016). A few PKs that function in response to biotic and abiotic stresses have been shown to undergo alternative splicing (Rostoks *et al*., 2004; Koo *et al*., 2007; Lin *et al*., 2010), which could explain the high frequency of this category with alternatively spliced PKs.

Overall, the Ssp and Sbi kinomes exhibited similar duplication patterns; in both species, the most common type of PK duplication was segmental duplication, followed by tandem duplications. These duplication events are usually reported as the two main contributors to PK expansion in the genomes of several other species, especially in the RLK-Pelle superfamily (Champion *et al*., 2004; Dardick *et al*., 2007; Wei *et al*., 2014; Liu *et al*., 2015; Dezhsetan, 2017; Zhu *et al*., 2018a, b). Gene retention by tandem duplication in kinases has already been identified, with very high rates in several plants (Lehti-Shiu and Shiu, 2012), and considerable correlation with different kinds of stress (Freeling, 2009). The association of PK expansion through such events with defense response and signaling pathways has been widely reported in kinome studies (Zulawski *et al*., 2014; Liu *et al*., 2015; Yan *et al*., 2018; Zhu *et al*., 2018a, b), with these events being more pronounced in the RLK-Pelle group. In the Ssp and Sbi kinomes, we found several subfamilies in this group with tandem duplications (mostly in LRR families). By analyzing GO biological process categories related to these events (Figs. 3 and 4), we found a considerable frequency of categories related to the defense response; however, other general categories were also frequent, which is explained by the numerous processes related to these subfamilies. Interestingly, in the RKF-3 family (in the RLK-Pelle group) in Ssp, all duplications were associated with tandem events, and members of this family have already been linked with stress responses and extracellular signaling (Huang *et al*., 2014; Vaid *et al*., 2016). Even with this high similarity, several differences in the distribution of tandemly organized genes within subfamilies were found between the Ssp and Sbi kinomes. These species- and chromosomal region-specific organizational characteristics were previously noted by Yan *et al*. (2018) in a comparison of cotton kinomes. With respect to genome organization in Ssp and Sbi, different forms of tandem events have already been found (Wang *et al*., 2010), with specific gene organization patterns within each genome (Zhang *et al*., 2018).

Segmental duplication events were also the major contributors to PK expansion in other species; in the soybean kinome, these events accounted for the origin of more than 70% of the PKs (Liu *et al*., 2015); in grapevine, they were estimated to be responsible for the origin of ∼30% of the kinases and were thought to be especially relevant in the expansion of the RLK-Pelle family (Zhu *et al*., 2018b). The most striking duplication-related difference between the Ssp and Sbi kinomes was the distribution of the rate of nonsynonymous mutations (Ks), which was used to estimate the time of occurrence of these segmental duplications. While the range of Sbi PK Ks values was comparatively wide, peaking at 0.65-0.85 (Fig. 2E), the Ks values of Ssp exhibited a very prominent peak between 0 and 0.05 range; in addition, the further distribution of Ks was somewhat similar to that in Sbi (Fig. 2F). Based on the clock-like rates of synonymous substitutions, we estimated that the time of occurrence of this large number of segmental duplications with Ks*<*0.05 was less than 3.8 MYA. Thus, we postulated that the Ks distribution in Ssp is a consequence of the recent polyploidization events in sugarcane; this hypothesis is supported by recent indications that the *Saccharum*-specific WGDs occurred in the last 3.1-4.6 million years (Kim *et al*., 2014; Zhang *et al*., 2018). This hypothesis is further reinforced by the findings reported in *Gossypium* spp. kinomes; a profile of Ks distributions very similar to that in Ssp was observed in the allotetraploids *G. hirsutum* and *G. barbadense* but not in its diploid relatives (Yan *et al*., 2018), strengthening the connection of this profile to WGD events.

We also analyzed the ratio of synonymous to nonsynonymous mutations (Ka/Ks), which is used to determine the type of selection acting on a gene (Zhang *et al*., 2006). We found that in the two kinomes the large majority of segmentally duplicated PKs were under negative selection (Ka/Ks*<*1), while a smaller percentage were under positive selection (Ka/Ks*>*1), and very few were under neutral selection (Ka/Ks=1). This pattern is similar to those observed in the soybean, grapevine and pineapple kinomes (Liu *et al*., 2015; Zhu *et al*., 2018a, b) and to those reported in smaller gene families in Ssp (Wang *et al*., 2019c; Li *et al*., 2020a) and sorghum (Malviya *et al*., 2016; Anand *et al*., 2017; Mittal *et al*., 2017).

Several RNA-Seq experiments were used to estimate the expression patterns of kinase subfamilies across a considerable range of tissues and genotypes. Due to the similar expression patterns within kinome subfamilies (Liu *et al*., 2015) and the possibility of detecting clearer expression patterns in different subfamilies than at the individual gene level, expression analysis was performed combining expression levels of genes from subfamilies, instead of individual genes. The differences among samples were more evident when separated by tissue instead of genotype and species (Fig. 7), possibly because of the strong conservation of PKs and their importance in several fundamental biological processes. In addition, as Liu *et al*. (2015) suggested, we also recognized that the Ssp and Sbi kinomes’ expression is shaped by the physiological characteristics of these species. The highest expression levels were found for members of the CMGC group in both the Sbi and Ssp kinomes. Additionally, in the AGC, CAMK, and CK1 groups, we found high expression levels in several subfamilies. These findings were previously reported in other plant species, suggesting an association of these groups with developmental processes (Liu *et al*., 2015; Zhu *et al*., 2018a, b). Interestingly, even though RLK-Pelle subfamilies account for the largest number of PKs among the kinomes, they were not among the top overexpressed subfamilies.

Among the 10% subfamilies with the highest expression, CMGC_GSK, CMGC_RCK, and CK1_CK1 were common to both the Sbi and Ssp datasets. Importantly, in addition to being overexpressed in Sbi, the members of the CMGC_GSK subfamily also contain many functional domains, as mentioned previously, and this characteristic might reflect their high expression. Another interesting subfamily with overexpression in the Sbi kinome was CMGC_MAPK (which exhibited average expression in Ssp), which has already been identified as having considerable importance in several Ssp and Sbi studies concerning stress signaling (Zhang *et al*., 2015; Paungfoo-Lonhienne *et al*., 2016; Li *et al*., 2016b; Srivastava and Kumar, 2020; Tuleski *et al*., 2020; Wang *et al*., 2020).

Despite having only one kinase member in the Ssp and Sbi kinomes, the AGC_PKA-PKG subfamily showed one of the highest average expression values across Ssp tissues. In addition to the unremarkable expression of RLK-Pelle subfamily members, the high AGC_PKA-PKG expression corroborates the observation that in the Ssp and Sbi kinomes, expression was not related to the number of family members across families and groups. If we assume that PK subfamilies may have increased in size through duplication events, this might be a case of dosage balance, a phenomenon wherein the function of regulatory genes is sensitive to a stoichiometric equilibrium (Birchler & Veitia, 2014). Thus, PK families composed of more members (which survived duplications and thus present more copies) have a tendency toward lower average expression, as has been demonstrated in other plants (Birchler & Veitia, 2012). Additionally, AGC_PKA-PKG subfamily has been reported as broadly important in both Ssp and Sbi. In Ssp, studies have demonstrated the association of its members with signaling pathways (Kasirajan *et al*., 2020), cell proliferation (Li *et al*., 2016b), infection responses (Santa Brigida *et al*., 2016; Xu *et al*., 2018), hormone signal transduction in response to drought (Li *et al*., 2016a), and pathways related to sucrose storage and photosynthesis (Hoang *et al*., 2017; Thirugnanasambandam *et al*., 2017). In Sbi, the importance of this family is also linked with stress and signal responses (Parra-Londono *et al*., 2018; Li *et al*., 2018b; Nagaraju *et al*., 2020; Vikal *et al*., 2020). Therefore, these insights into expression constitute a valuable reservoir of information for analyzing the importance of Ssp and Sbi kinases.

The final analysis performed using RNA-Seq datasets aimed to establish closer relationships among kinase subfamilies in Sbi and Ssp through coexpression networks, enabling biological inferences using connection patterns. The gene coexpression networks were constructed with pairwise correlations (similarity scores) from the gene expression quantification data (Serin *et al*., 2016). Pearson correlation coefficients were used because of their reasonable performance in RNA-Seq datasets (Ballouz *et al*., 2015). Moreover, as Liu *et al*. (2015) suggested, we constructed the networks based on subfamily relationships instead of single genes because of the enhanced functional interpretability and general inferences allowed by this approach. Complex networks have been widely applied to visualize complex biological systems (Barabási, 2016), and constitute a powerful tool for modeling gene interactions (Zhao *et al*., 2010). For kinase subfamily representations, these networks can facilitate the interpretation of relevant relationships among sets of kinases and provide insights into the interactions among metabolic mechanisms. These applications are possible because similar expression patterns on genes belonging to the same pathways reflect the network structure (Lee *et al*., 2015), thus providing a tool to model these complex molecular interactions (Ficklin and Feltus, 2011).

Together with the network representations, we used community detection methodologies to identify modules of cohesive elements, which possibly indicate more strongly interconnected metabolic relationships (Mitra *et al*., 2013; Mall *et al*., 2017; Zhang and Yin, 2020). This structural organization constitutes a reservoir of genetic information among kinomes and provides important insights into how these subfamilies biologically interact. When some subfamilies without significant relationships with other elements (disconnected nodes) were excluded, the network structures (Fig. 8) indicated that all of the subfamilies were interconnected, considering the nonrandom dependencies across subfamilies captured by the established correlation coefficient threshold (Ficklin and Feltus, 2011). This connected structure observed among both the Sbi PKs and the Ssp PKs has been described throughout the manuscript. Even though they have specific functions, all kinase subfamilies play roles in several common metabolic processes, and this commonality is clearly reflected in the network structures.

In addition, considering the roles of PKs in metabolic signaling and stress responses, the organization of several subfamilies is reasonably conserved among different plant species (Lehti-Shiu and Shiu, 2012). By comparing the Sbi and Ssp networks, we identified a substantial core of similarity between the subfamily interactions in these species, possibly indicating several analogous expression profiles, as already observed in the comparison of expression values among tissues and genotypes (Fig. 7). In addition to the comparison of network connectivities, other topological characteristics were used to identify important features in the organization of kinome subfamilies. Hub and betweenness measures were calculated to supply evidence regarding how specific subfamilies are important in most metabolic processes involving kinases.

Within a network structure, elements with the most connections are called hubs (Barabási, 2016). These nodes have been used to identify functionally critical components and as an additional approach to describe the network structure (Hong *et al*., 2013; Azuaje, 2014; Van Dam *et al*., 2018). In the constructed networks, the hub nodes indicate kinase subfamilies with the most correlations, which might represent sets of kinases with influential roles in diverse metabolic mechanisms in kinomes. Interestingly, the Sbi and Ssp networks did not exhibit high overlap of hub nodes. This observation provides evidence that although there are several similarities among the kinase expression profiles in these species, and the same biological cascades are activated, as indicated by the GO analyses (Supplementary Fig. S5), the mechanism by which the expression balance is achieved is species-specific. In fact, previous studies in polyploids have shown that this balancing varies even among lines of the same species (Mutti *et al*., 2017).

In both these Sbi and Ssp networks and those constructed in other kinome studies (Liu *et al*., 2015; Zhu *et al*., 2018b), different members of the RLK-Pelle group (mostly those in the LRR and RLCK families) were identified as hubs. Considering the described abundance of these families, their tandem duplications, and related functional implications, the strong influence of such nodes on the correlations among kinase subfamilies was expected. CMGC group subfamilies were also identified as hub elements, as observed in the soybean kinome (Liu *et al*., 2015). In the sugarcane network, the GSK and CDK families had a considerable number of connections, which is clearly explained by the very high number of pathways in which their members are involved, as previously noted. Additionally, the CDK family has already been found to be related with stress signaling in Ssp (Patade *et al*., 2011) and in Sbi (Challa and Neelapu, 2018). In Sbi, the DYRK family also had a high node degree. Interestingly, members of this family have already been found to be related to the suppression of photosynthesis activity (Kimura and Ishikawa, 2018); thus, the importance and impact of this family among kinases is evident. Among the other hubs, the STE group (STE20 family) was also important in the Ssp and Sbi networks, which can be explained by the high number of biological cascades related to this subfamily (Xiong *et al*., 2016).

Several factors can explain why a subfamily constitutes a hub in our constructed networks, such as a high expression level (CMGC_GSK, CAMK_OST1L, CK1_CK1, and CMGC_DYRK-YAK), a large number of subfamily members (RLK-Pelle_LRR-III), the occurrence of tandem duplications (IRE1), a more structured gene organization considering intron-exon structures (RLK-Pelle_RLCK-XII-1, RLK-Pelle_LRR-VI-1, and CMGC_GSK), and the presence of diverse functional domains (RLK-Pelle_LRR-III, CMGC_CDK-CRK7-CDK9, and RLK-Pelle_LRR-VII-1) or multikinase domains (AGC_RSK-2, RLK-Pelle_LRR-III, and CMGC_CDK-CRK7-CDK9). However, we did not observe a consistent feature profile required for a subfamily to be considered a hub. Evidence supports the hubs’ importance; however, the real reasons for their key importance within these structures are likely to be linked with functional properties, as widely discussed in other coexpression studies (Goel *et al*., 2018; Tai *et al*., 2018; Wang *et al*., 2018; Zou *et al*., 2019; Ding *et al*., 2020).

In addition to hub descriptions, edge betweenness measures also have high interpretability considering the complex system modeled by the networks. These calculations are based on properties from the entire network (Dunn *et al*., 2005), exploiting the network flow and identifying possible essential interactions for the visual configuration (Van Dam *et al*., 2018). In both the Sbi and Ssp networks, a clearly separated group of PKs that was connected with the other elements by only one or a few connections was evident. This network configuration might indicate important relationships among kinase subfamilies, providing evidence indicating how these specific subfamilies can interconnect. In Ssp, the most critical connections identified by betweenness calculations were found in the RLK-Pelle_L-LEC/RLK-Pelle_LRR-VIII-1 and RLK-Pelle_CR4L/RLK-Pelle_LRR-Xb-1 subfamilies. These nodes are members of families with undeniable importance, as seen in the network structure. The bridges in these kinase-kinase interactions can be explained by the large number of members within these families that can act in a connected manner, which is less evident in other subfamilies. However, as observed in the hub configurations, these structures are more evidently linked with functional roles, such as interconnected signaling pathways.

In the Sbi network, on the other hand, the highest betweenness values were found in the CAMK_CDPK/RLK-Pelle_LRR-VI-2 and CAMK_CDPK/CMGC_RCK subfamilies. Interestingly, CAMK_CDPK subfamily genes have been extensively indicated to be located at important genomic regions regulating plant growth, development and resistance mechanisms to several types of abiotic and biotic stresses in both Sbi (Pestenácz and Erdei, 1996; Nhiri *et al*., 1998; Jain *et al*., 2008; Li *et al*., 2010; Monreal *et al*., 2013; Usha Kiranmayee *et al*., 2017) and Ssp (Li *et al*., 2016b; Marquardt *et al*., 2017; Ling *et al*., 2018; Dharshini *et al*., 2020; Srivastava and Kumar, 2020; Su *et al*., 2020b), further supporting the association of functional characteristics in the network structure.

## 5. Conclusions

This study provided an extensive reservoir of genetic and molecular information for both Sbi and Ssp. Considering the incontestable importance of kinases in several essential biological processes, the identification, categorization and analysis of the kinomes of these species resulted in an important compendium of knowledge for use in further studies. Clear similarities were found in protein properties, domain compositions, genomic organization, expression profiles and subfamily interactions. However, we also observed pronounced differences in duplication events, which probably arose from Ssp recent WGDs, facilitating understanding of how the Sbi and Ssp kinomes have evolved considering this vast protein superfamily.

## Supplementary data

### Supplementary figures

***Fig. S1.*** (Additional File 1) Phylogenetic analysis of the identified protein kinases in *Sorghum bicolor* (Sbi) with 1,000 bootstrap replicates. Each protein is separated on the right side of the tree and is presented with its classification with respect to the kinase subfamilies, which are colored to represent the differences among subfamilies.

***Fig. S2*.** (Additional File 2) Phylogenetic analysis of the identified protein kinases in *Saccharum spontaneum* (Ssp) with 1,000 bootstrap replicates. Each protein is separated on the right side of the tree and presented with its classification with respect to the kinase subfamilies, which are colored to represent the differences among subfamilies.

***Fig. S3*.** (Additional File 3) Phylogenetic analysis of the protein kinases identified in both *Sorghum bicolor* (Sbi) and *Saccharum spontaneum* (Ssp) with 1,000 bootstrap replicates. Each protein is separated on the right side of the tree and presented with its classification with respect to the kinase subfamilies, which are colored to represent the differences among subfamilies.

***Fig. S4.*** (Additional File 4) Kinase subfamily quantification analysis in different plant species. Each row indicates a different subfamily and each column a plant species, and the numbers of kinases are noted. This heatmap is colored according to the distribution of quantities present in the datasets on a scale of beige to dark green.

***Fig. S5*.** (Additional File 5) Gene Ontology (GO) category annotation of biological processes in (A) the entire set of *Saccharum spontaneum* (Ssp) kinases; (B) the entire set of *Sorghum bicolor* (Sbi) kinases; and (C) the set of Sbi kinases related to alternative splicing events.

***Fig. S6.*** (Additional File 6) Venn diagram showing the intersection of subfamilies in the communities within the *Sorghum bicolor* (Sbi) and *Saccharum spontaneum* (Ssp) networks.

***Fig. S7.*** (Additional File 7) Coexpression network for *Sorghum bicolor* (Sbi) kinase subfamilies. Each node corresponds to a different subfamily, its size corresponds to the average expression value for all kinases within the subfamily in different samples, and its color corresponds to the hub score and ranges from beige to dark green. Each edge corresponds to a correlation with a Pearson correlation coefficient of at least 0.6. The correlation strength is represented by the edge’s width, and the edge betweenness score is represented by the color (ranging from black to light blue, with light blue representing the highest values). The network background is colored according to the community detection analysis, and the nodes are labeled according to the subfamily correspondence found in Supplementary Table S40.

***Fig. S8*.** (Additional File 8) Coexpression network for *Saccharum spontaneum* (Ssp) kinase subfamilies. Each node corresponds to a different subfamily, its size corresponds to the average expression value of all kinases within the subfamily in different samples, and its color corresponds to the hub score and ranges from beige to dark green. Each edge corresponds to a correlation with a Pearson correlation coefficient of at least 0.6. The correlation strength is represented by the edge’s width, and the edge betweenness score is represented by the color (ranging from black to light blue, with light blue representing the highest values). The network background is colored according to the community detection analysis, and the nodes are labeled according to the subfamily correspondence found in Supplementary Table S41.

### Supplementary tables

#### Additional file 9

***Table S1.*** Organization of sorghum RNA-Seq experiments.

***Table S2.*** Organization of sugarcane RNA-Seq experiments.

***Table S3.*** Kinase domain annotation of the 1,210 sorghum protein kinases.

***Table S4.*** Kinase domain annotation of the 2,919 sugarcane protein kinases.

***Table S5.*** Subfamily kinase classification of the sorghum 1,210 kinases based on the alignment on HMMER and confirmed by phylogeny.

***Table S6.*** Subfamily kinase classification of the sugarcane 2,919 kinases based on the alignment on HMMER and confirmed by phylogeny.

***Table S7.*** Sorghum and sugarcane kinase subfamily quantifications.

***Table S8.*** Sorghum kinase distribution across chromosomes.

***Table S9.*** Sugarcane kinase distribution across chromosomes and alleles.

***Table S10.*** Localization, intron quantity and possible alternative splicing events of the 1,210 sorghum kinases.

***Table S11.*** Localization and intron quantity of the 2,919 sugarcane kinases.

***Table S12.*** Domain annotation of the 1,210 sorghum protein kinases.

***Table S13.*** Domain organization of the 1,210 sorghum protein kinases.

***Table S14.*** Sorghum kinase domain organization for proteins with multiple kinase domains.

***Table S15.*** Domain annotation of the 2,919 sugarcane protein kinases.

***Table S16.*** Domain organization of the 2,919 sugarcane protein kinases.

***Table S17.*** Sugarcane kinase domain organization for proteins with multiple kinase domains.

***Table S18.*** Gene Ontology (GO) annotations for the 1,210 sorghum kinases.

***Table S19.*** Gene Ontology (GO) annotations for the 2,919 sugarcane kinases.

***Table S20.*** Compositional analyses of the 1,210 sorghum kinases.

***Table S21.*** Compositional analyses of the 2,919 sugarcane kinases.

***Table S22.*** Characteristics of sorghum kinase subfamilies.

***Table S23.*** Characteristics of sugarcane kinase subfamilies.

***Table S24.*** Presence of domains across sorghum kinase subfamilies.

***Table S25.*** Presence of domains across sugarcane kinase subfamilies.

***Table S26.*** Duplication origin of the 1,210 sorghum kinases.

***Table S27.*** Duplication origin of the 2,919 sugarcane kinases.

***Table S28.*** Collinearity events and Ka/Ks values of sorghum protein kinases.

***Table S29.*** Collinearity events and Ka/Ks values of sugarcane protein kinases.

#### Additional file 10

***Table S30.*** Sorghum kinase transcripts per million (TPM) values across samples.

***Table S32.*** Sorghum kinase subfamily quantification across samples.

***Table S34.*** Sorghum kinase subfamily quantification across tissues from the selected genotypes.

***Table S36.*** Descriptive statistics of subfamily expression across sorghum kinase subfamilies.

***Table S38.*** Spearman correlation of average transcripts per million (TPM) values in sorghum genotypes/tissues with kinase subfamily quantities.

***Table S40.*** Sorghum kinase subfamily coexpression network characterization.

***Table S42.*** Edge betweenness values calculated across the sorghum coexpression network.

#### Additional file 11

***Table S31.*** Sugarcane kinase transcripts per million (TPM) values across samples.

***Table S33.*** Sugarcane kinase subfamily quantification across samples.

***Table S35.*** Sugarcane kinase subfamily quantification across tissues from the selected geno-types.

***Table S37.*** Descriptive statistics of subfamily expression across sugarcane kinase subfamilies.

***Table S39.*** Spearman correlation of average transcripts per million (TPM) values in sugarcane genotypes/tissues with kinase subfamily quantities.

***Table S41.*** Characterization of the sugarcane kinase subfamily coexpression network.

***Table S43.*** Edge betweenness values calculated across the sugarcane coexpression network.

#### Additional file 12

***Table S44.*** Sugarcane RNA-Seq read counts considering the samples described in Supplementary Table S2 and Saccharum spontaneum coding DNA sequences.

#### Additional file 13

***Table S45.*** Sugarcane RNA-Seq transcripts per million (TPM) values considering the samples described in Supplementary Table S2 and Saccharum spontaneum coding DNA sequences.

## Acknowledgements

This work was supported by grants from the Fundação de Amparo à Pesquisa de do Estado de São Paulo (FAPESP 2020/07434-0, 2015/22993-7, 2008/52197-4 and 2005/55258-6), the Conselho Nacional de Desenvolvimento Científico e Tecnológico (CNPq 313426/2018-0 and 434886/2018-1), and the Coordenação de Aperfeiçoamento de Pessoal de Nível Superior (CAPES—Computational Biology Programme and Financial Code 001). AHA received a PhD fellowship from FAPESP (2019/03232-6); RJGP received MSc fellowships from CAPES (88887.177386/2018-00) and from FAPESP (2018/18588-8); GKH received a PhD fellowship from CAPES (88882.160212/2017-01); CBCS received a PD fellowship from FAPESP (2015/16399-5); MCM received a PD fellowship from FAPESP (2014/11482-9); DAS received a PhD fellowship from FAPESP (2010/50119-6); LBS received an undergraduate fellowship from FAPESP (2019/19340-2); TWB received a PhD fellowship from FAPESP (2010/50091-4); and APS received a research fellowship from CNPq.

## Author contributions

AHA and RJGP performed all analyses and wrote the manuscript. ALBG, FHC, and GKH assisted in processing the sugarcane RNA-Seq data, and together with CBCS, MCM, and DAS, were responsible for the sugarcane RNA-Seq experiments. MMC assisted in the functional analyses of the kinase subfamilies; LBS contributed to the identification of the kinases; JSN assisted in the kinase categorization; LRP, MGAL, MSC, and TWB were responsible for the sugarcane field experiments; MGQ assisted in the network analysis; WAP assisted in the pipeline definition and kinase categorization; and GRAM and APS conceived the project. All authors reviewed, read, and approved the manuscript.

## Data availability

The data that support the findings of this study are openly available in https://doi.org/10.6084/m9.figshare.c.5122460.v1, as well supplementary data.

## Abbreviations

Aco: Aquilegia coerulea
AGC: cyclic AMP-dependent protein kinase (cAPK), cGMP-dependent protein kinase, and lipid signaling kinase families
Aly: Arabidopsis lyrata
Ath: Arabidopsis thaliana
B-lectin: D-mannose-binding lectin
Bdi: Brachypodium distachyon
CAMK: calcium- and calmodulin-regulated kinase
Ccl: Citrus clementina
CDS: DNA coding sequence
CK1: casein kinase 1
CMGC: cyclin-dependent kinase, mitogen-activated protein kinase, glycogen synthase kinase and cyclin-dependent kinase-like kinase
Cpa: Carica papaya
Cre: Chlamydomonas reinhardtii
Csa: Cucumis sativus
Csi: Citrus sinensis
DUF26: Domain of Unknown Function 26
Egr: Eucalyptus grandis
ER: endoplasmic reticulum
GM: gene model
Gma: Glycine max
GO: Gene Ontology
GUB: galacturonan-binding
HMM: hidden Markov model
IRE1: inositol-requiring kinase 1
Ka: Nonsynonymous substitution rates
Ks: Synonymous substitution rates
LRR: leucine-rich repeat
LRRNT: leucine-rich repeat N-terminal domain
Mes: Manihot esculenta
Mgu: Mimulus guttatus
Mtr: Medicago truncatula
MW: molecular weight
MYA: million years ago
Osa: Oryza sativa
PEK: pancreatic eukaryotic initiation factor-2alpha kinase
PK: protein kinase
pI: isoelectric point
Ppa: Physcomitrella patens
Ppe: Prunus persica
Ptr: Populus trichocarpa
Rco: Ricinus communis
RLK: receptor-like kinase
S-locus-glycop: S-locus glycoprotein
Sbi: Sorghum bicolor
Sit: Setaria italica
Smo: Selaginella moellendorffii
Ssp: Saccharum spontaneum
STE: serine/threonine kinase
TKL: tyrosine kinase-like kinase
TPM: Transcripts per million
Vca: Volvox carteri
Vvi: Vitis vinifera
WAK: wall-associated receptor kinase
WGD: whole-genome duplication
Zma: Zea mays

## Notes

https://doi.org/10.6084/m9.figshare.c.5122460.v1

### Competing Interest Statement

The authors have declared no competing interest.

